# Distinct subcellular localizations of DUF1218 proteins in *Marchantia polymorpha* and *Nicotiana benthamiana* reveal two different plasmodesmata-targeting mechanisms

**DOI:** 10.64898/2026.08.17.745377

**Authors:** Linh Ta Thi Thuy, Shiuan-Jie Tsai, Sumanth K. Mutte, Hsiang-Chi Lee, Chia-Ming Hsu, Hui-Yu Chang, Kuan-Ju Lu

## Abstract

Plasmodesmata are membrane-lined channels connecting plant cells to facilitate intercellular transport of molecules. Although many plasmodesmata-localized proteins have evolved throughout plant evolution, whether they use a conserved targeting system remains unclear. In the bryophyte *Marchantia polymorpha*, we identified two DUF1218-domain proteins homologous to the Arabidopsis plasmodesmata-localized AtTVA. When ectopically expressed, MpDUF1218-1 localized to plasmodesmata in both *Nicotiana benthamiana* and *M. polymorpha*, whereas MpDUF1218-2 formed cytoplasmic puncta in both species. Unexpectedly, AtTVA formed cytoplasmic puncta rather than localizing to plasmodesmata in *M. polymorpha*. Domain-swap analyses revealed that the first helix of MpDUF1218-1 is crucial for plasmodesmata localization in *N. benthamiana*, while the first two helices are required in *M. polymorpha*. In contrast, the second and third helices of AtTVA contribute to its plasmodesmata localization in *N. benthamiana*. Further domain dissection indicated that other regions of MpDUF1218-1 also contribute to accurate targeting by regulating its distribution among the ER, cytoplasmic puncta, and plasma membrane. Together, our findings suggest that MpDUF1218-1 is targeted by a mechanism shared between the two species, whereas AtTVA relies on a distinct mechanism present in *N. benthamiana* but absent in *M. polymorpha*, suggesting the emergence of alternative plasmodesmata-targeting pathways during land plant evolution.

**Highlight:** Comparative analysis of DUF1218 proteins reveals the diversification of plasmodesmata-targeting mechanisms, providing insights into the evolution of intercellular communication in land plants.

## Introduction

Cell-to-cell communication is crucial for the development of multicellularity. A major and unique route for cell-to-cell signaling in plants is mediated by cell wall-embedded channels termed plasmodesmata (Bayer and Benitez-Alfonso, 2024). Plasmodesmata are composed of a plasma membrane-lined cylindrical pore with a compressed central desmotubule derived from the endoplasmic reticulum. Spaces between the plasma membrane and the desmotubule are called cytoplasmic sleeves, which provide the primary route for molecules to travel between cells (Kragler *et al*., 1998, Brunkard and Zambryski, 2016, Sager and Lee, 2018, Li *et al*., 2021, Bayer and Benitez-Alfonso, 2024). Plasmodesmata not only allow the transport of small molecules such as nutrients and hormones, but also macromolecules, including regulatory proteins, DNA/RNA molecules, and even some organelles and plant viruses may spread from cell to cell via plasmodesmata (Liu and Chen, 2018, Lu *et al*., 2018, Gundu *et al*., 2020, Reagan and Burch-Smith, 2020, Kumar and Dasgupta, 2021, Li *et al*., 2021, Tee and Faulkner, 2024) Plasmodesmata are crucial for the survival of all land plants as is reflected in their presence in all land plants (Raven, 2007, Brunkard and Zambryski, 2017, Tee and Faulkner, 2024). Nevertheless, during the evolution of land plants, plasmodesmata might have evolved different properties to serve different purposes. In Arabidopsis, different forms of plasmodesmata have been discovered (Bayer and Benitez-Alfonso, 2024). Recently, one of the specialized plasmodesmata, named funnel-shaped plasmodesmata, was identified to facilitate the bulk transportation of molecules between protophloem sieve elements and phloem pore pericycle cells (Ross-Elliott *et al*., 2017). However, the capacity to generate complex plasmodesmata seems to be lost in one of the bryophytes, *Physicometrium patens* during the evolution while the liverwort, *Marchantia polymorpha* and the hornwort, *Anthoceros agrestis* can still generate complex plasmodesmata (Wegner and Ehlers, 2024, Hsu *et al*., 2025). The diversification of the ability to build different plasmodesmata also demonstrates that plasmodesmata may be modified during evolution.

In addition to the variety of structures, the molecular composition of plasmodesmata may also diverge during evolution. The Arabidopsis plasmodesmata proteome revealed many structural proteins as well as those involved in the regulation of intercellular transport (Fernandez-Calvino *et al*., 2011). One such protein family is PLASMODESMATA-LOCALIZED PROTEINS (PDLPs). PDLPs are known to be localized at plasmodesmata and regulate the accumulation of callose (β-1,3-glucan) around the neck region of plasmodesmata to modulate their permeability (Thomas *et al*., 2008, Lee *et al*., 2011). Notably, PDLP proteins evolved after the emergence of vascular tissue in land plants (Vaattovaara *et al*., 2019), suggesting that the molecular composition of plasmodesmata may have changed along evolution.

For plant cells, assigning newly evolved proteins to plasmodesmata means adaptation of the existing sorting system. Alternatively, a new mechanism might evolve to accommodate new proteins. It has been demonstrated that the movement protein of tobacco mosaic virus (TMVMP) targets preferentially the complex plasmodesmata (Ding *et al*., 1992), and TMVMP does not colocalize with the known plasmodesmata protein, PDLP5 (Lee *et al*., 2011). Additionally, the targeting to plasmodesmata of another PDLP, PDLP1a from *Nicotiana benthamiana*, is not affected by the treatment of Brefeldin A (BFA), which is known to block the targeting of TMVMP to plasmodesmata (Tagami and Watanabe, 2007, Wright *et al*., 2007, Thomas *et al*., 2008). These results imply that plant cells might employ specific mechanisms to direct different cargos to different types of plasmodesmata. However, how do these mechanisms evolve along the evolution is still not clear.

In this research, we investigated whether plasmodesmata-targeting mechanisms are conserved between plants from different linages. We isolated homologs of known plasmodesmata proteins from *M. polymorpha* and tested their subcellular localization in the vascular plant *N. benthamiana*. We identified two homologs of Arabidopsis AtTVA (TRANVIA, At3G15480, a DUF1218 domain-containing protein) (Fernandez-Calvino *et al*., 2011) in *M. polymorpha* and found that MpDUF1218-1 (Mp8g06520) localized to plasmodesmata in *N. benthamiana* while MpDUF1218-2 (Mp8g17930) formed puncta at cell periphery but did not locate to plasmodesmata. In *M. polymorpha*, both MpDUF1218-1 and -2 showed similar subcellular localization as in *N. benthamiana*. However, AtTVA did not locate to plasmodesmata, which was distinct from its localization in Arabidopsis and *N. benthamiana*. Phylogenetic analysis revealed AtTVA-clade DUF1218 domain proteins branched after the emergence of Gymnosperms, suggesting an evolution event may have driven their divergence. Through a series of domain-swap and domain-deletion analyses, our study revealed that the *M. polymorpha* has a plasmodesmata-targeting mechanism which recognize MpDUF1218-1 while *N. benthamiana* has two systems, one recognizes MpDUF1218-1 and the other recognizes AtTVA. These results suggest that a complex system is evolved to fine-tune the targeting of plasmodesmata proteins in different plant species.

## Results

### In *Nicotiana benthamiana,* only one of the two MpDUF1218 proteins is localized to plasmodesmata

To investigate whether plasmodesmata-targeting mechanisms are conserved between plants from different linages, we performed a cross-species expression assay to observe the localization of proteins. By using the Arabidopsis plasmodesmata proteins AtREM1.3 (Remorin 1.3, AT2g45820), AtTET3 (TETRASPANIN 3, At3g45600), and AtTVA (TRANVIA, At3g15480) (Fernandez-Calvino *et al*., 2011) as the query proteins to BLAST against the *M. polymorpha* database (Marpol base, https://marchantia.info/) (Kawamura *et al*., 2022), we identified three proteins containing the Remorin C-terminal domain (Mp2g06670, Mp2g17160 and Mp2g28990), two tetraspanin proteins, MpTET3a (Mp3g07130) and MpTET3b (Mp6g10130), and four DUF1218-containing proteins: MpDUF1218-1 (Mp8g06520), MpDUF1218-2 (Mp8g17930), MpDUF1218-3 (Mp8g05730), and MpDUF1218-4 (Mp3G02810). A significant drop in the E-value comparing MpDUF1218-1, and -2 with MpDUF1218-3, and -4 led to the decision to focus on MpDUF1218-1 and -2 for further analysis (Fig. S1). We fused all homologs we cloned with fluorescent protein mGFP5 or mCitrine at the C-terminal and transiently expressed all fusion proteins by the constitutive 35S promoter in *N. benthamiana* leaf epidermal cells by agro-infiltration. We used aniline blue staining to label the callose that accumulates at plasmodesmata as a marker for plasmodesmata (Fig. 1 and 2).

**Figure 1.**
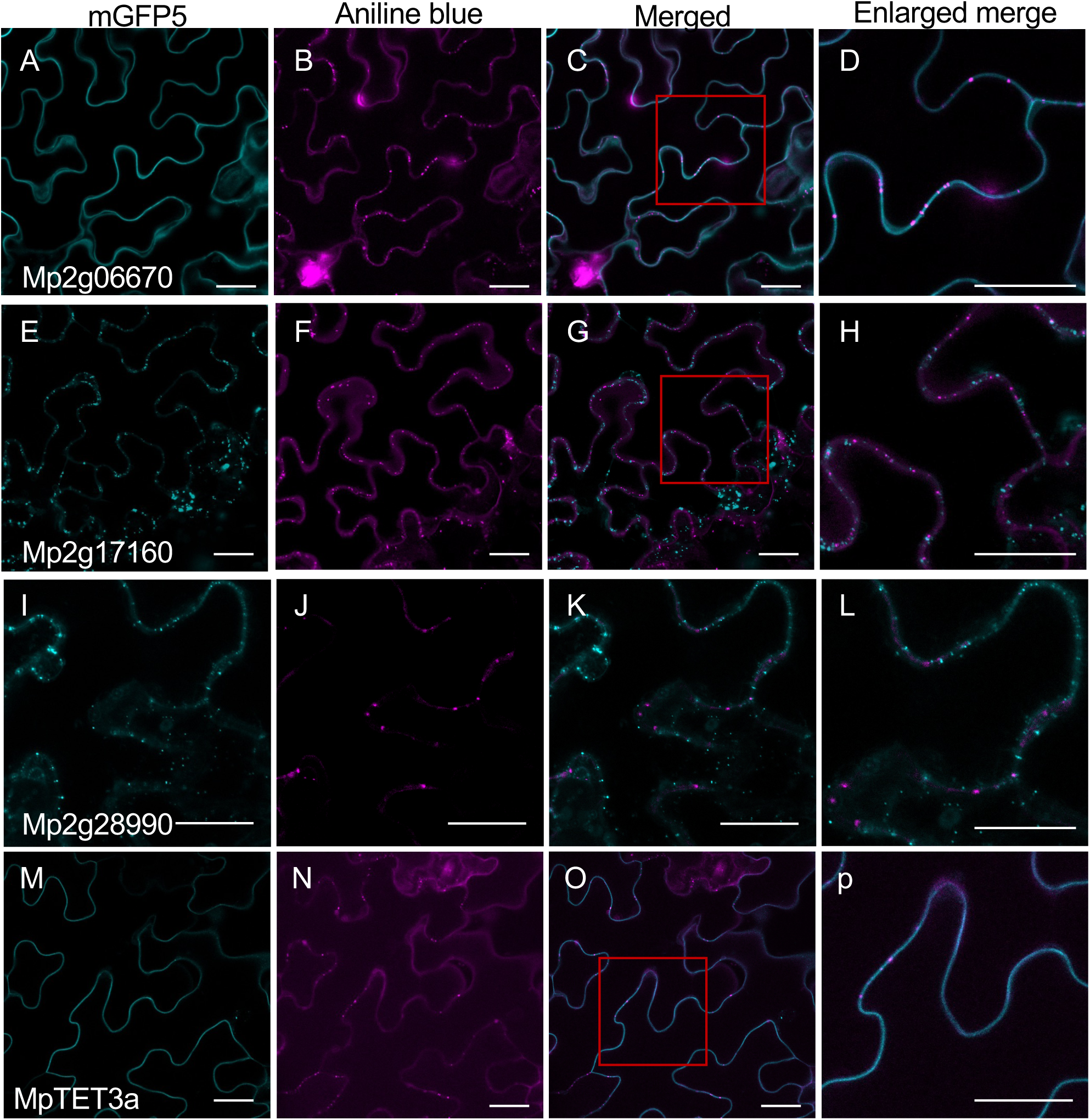
Colocalization of potential *Marchantia polymorpha* PD-localized proteins with aniline blue staining in *Nicotiana benthamiana*. (A, E, I, M) Fluorescent signals of the indicated fusion proteins expressed in epidermal cells of *N. benthamiana*. (B, F, J, N) Plasmodesmata were visualized using aniline blue staining on the same leaf sections. (C, G, K, O) Merged images of the first two panels, respectively. (D, H, and L) 2.5-fold and 1.5-fold (L) enlargement of the red boxed sections in C, G, O, and K, respectively. Scale bar, 30 μm.

**Figure 2.**
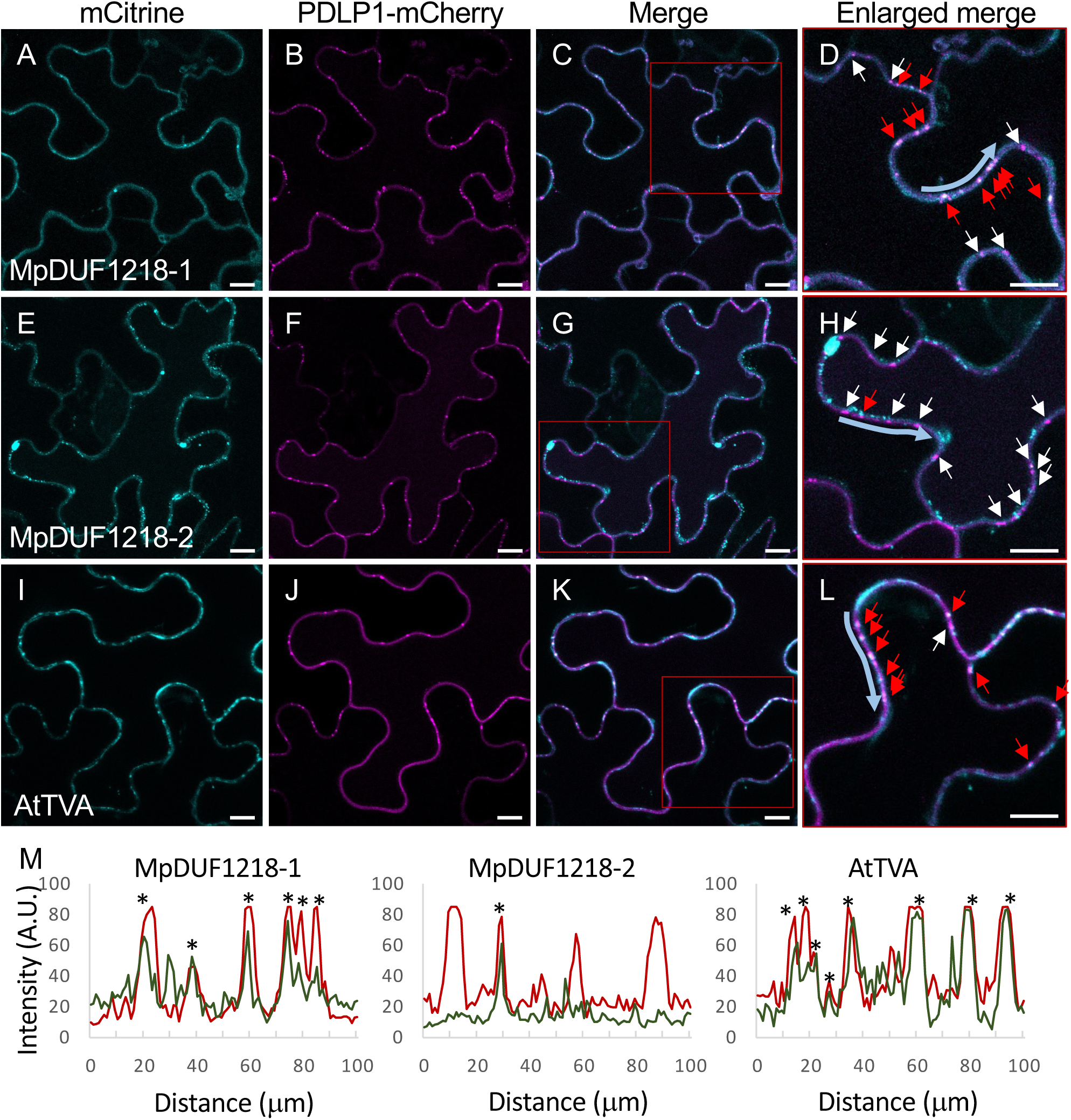
Colocalization of p*35S*::MpDUF1218-1-, MpDUF1218-2, and AtTVA-mCitrine with p*35S*::PDLP1-mCherry in *Nicotiana benthamiana*. (A, E, I) Fluorescent signals of the indicated fusion proteins expressed in epidermal cells of *N. benthamiana*. (B, F, J) Plasmodesmata were visualized using p*35S*::PDLP1-mCherry on the same leaf sections. (C, G, K) Merged images of the first two panels, respectively. (D, H, L) Two-fold enlargement of the red boxed sections in C, G, and K. Red arrows point to colocalization of the candidate proteins with the plasmodesmata markers. White arrows point to plasmodesmata marker only. Scale bar, 10 μm. (M) Colocalization plots show intensity profiles of the mCitrine fusion proteins (green lines) and PDLP1-mCherry (red lines) measured along the cell walls indicate by the blue arrows in D, H, and L. The x-axis is the distance along the cell wall in mm; the y-axis is the mean intensity of green and red channels in arbitrary units (A. U.). Asterisks mark the positions of overlapping peaks.

If a conserved plasmodesmata-targeting mechanism were present, we expected some homologs from these families would already acquire plasmodesmata localization properties and localized to plasmodesmata in *N. benthamiana*. In the Remorin family, we observed that Mp2g06670-mGFP5 distributed ubiquitously at the plasma membrane with no obvious enrichment (Fig. 1A). Mp2g17160- and Mp2g28990-mGFP5 formed puncta on the membrane but did not colocalize with the aniline blue signals (Fig. 1G, H, K and L). Similarly, the MpTET3a-mGFP5 located evenly on the plasma membrane and formed few enriched spots which did not colocalized with aniline blue signals (Fig. 1O and P). In addition, we could not clone MpTET3b, potentially due to its limited expression in the sexual organs as the annotation at Marpol base.

When we expressed *M. polymorpha* DUF1218 family proteins ectopically in *N. benthamiana*, we observed that both MpDUF1218-1 and -2 formed puncta structures at cell peripheries (Fig 2A, E). To test whether the puncta overlapped with plasmodesmata, we co-expressed the PLASMODESMATA-LOCALIZED PROTEIN 1-mCherry (PDLP1-mCherry) (Thomas *et al*., 2008) with all the DUF1218 proteins. The adaptation of PDLP1-mCherry was to avoid the physical perturbation during aniline blue staining process and we also verified PDLP1-mCherry as a reliable plasmodesmata marker by aniline blue staining (Fig. S2). We observed overlapping signals between PDLP1-mCherry and MpDUF1218-1-mCitrine (Fig. 2C, D). On the contrary, the MpDUF1218-2-mCitrine puncta did not colocalize with PDLP1-mCherry (Fig. 2G, H). These results suggest that MpDUF1218-1 can be recognized by the plasmodesmata-targeting system of *N. benthamiana* but not MpDUF1218-2. We also cloned and expressed AtTVA-mCitrine to confirm its plasmodesmata localization. Consistent with the published work, AtTVA-mCitrine formed enriched puncta at cell periphery and these puncta also colocalized with PDLP1-mcherry (Fig. 2I-L) (Fernandez-Calvino *et al*., 2011). We also measured the signal intensity of mCitrine and mCherry along the indicated length of cell membranes (Fig. D, H, L). The displayed plots demonstrate that most peaks of MpDUF1218-1- and AtTVA-mCitrine overlapped with the peaks of PDLP1-mcherry, which was not the case for MpDUF1218-2-mCitrine (Fig. 2M). Taken together, we suggest that both AtTVA and MpDUF1218-1 contain intrinsic properties that can be recognized by the plasmodesmata-targeting machinery in *N. benthamiana*, while MpDUF1218-2 does not contain such properties. Due to the DUF1218 family proteins were the only plasmodesmata-localized family amount our selected ones, we decided to further explore the evolution of the DUF1218’s plasmodesmata-targeting mechanisms.

### In *Marchantia polymorpha*, MpDUF1218-1 is localized to plasmodesmata but AtTVA and MpDUF1218-2 are not

We next investigate whether MpDUF1218-1, MpDUF1218-2, and AtTVA also localize to plasmodesmata in *M. polymorpha.* We generated p*35S*::MpDUF1218-1-, MpDUF1218-2-, and AtTVA-mCitrine transgenic *M. polymorpha* plants and investigated the expression pattern in dormant gemmae picked directly from the gemma cups (Fig. 3). Although at various degrees, the cell outline was visible in both MpDUF1218-1- and AtTVA-mCitrine gemmae but was barely distinguishable in MpDUF1218-2-mCitrine gemmae (Fig. 3B, E, H), indicating that the MpDUF1218-2 protein did not localize to the membrane (Fig. 3D-F). All three proteins accumulated as puncta inside epidermal cells (Fig. 3B, E, H) but only in MpDUF1218-1-mCitrine gemmae did we observe paired fluorescent puncta along the cell membrane, a pattern typical of plasmodesmata (Fig. 3C, red arrows). Although AtTVA-mCitrine signals were found at the cell membrane, no plasmodesmata-like localizations were found (Fig. 3I).

**Figure 3.**
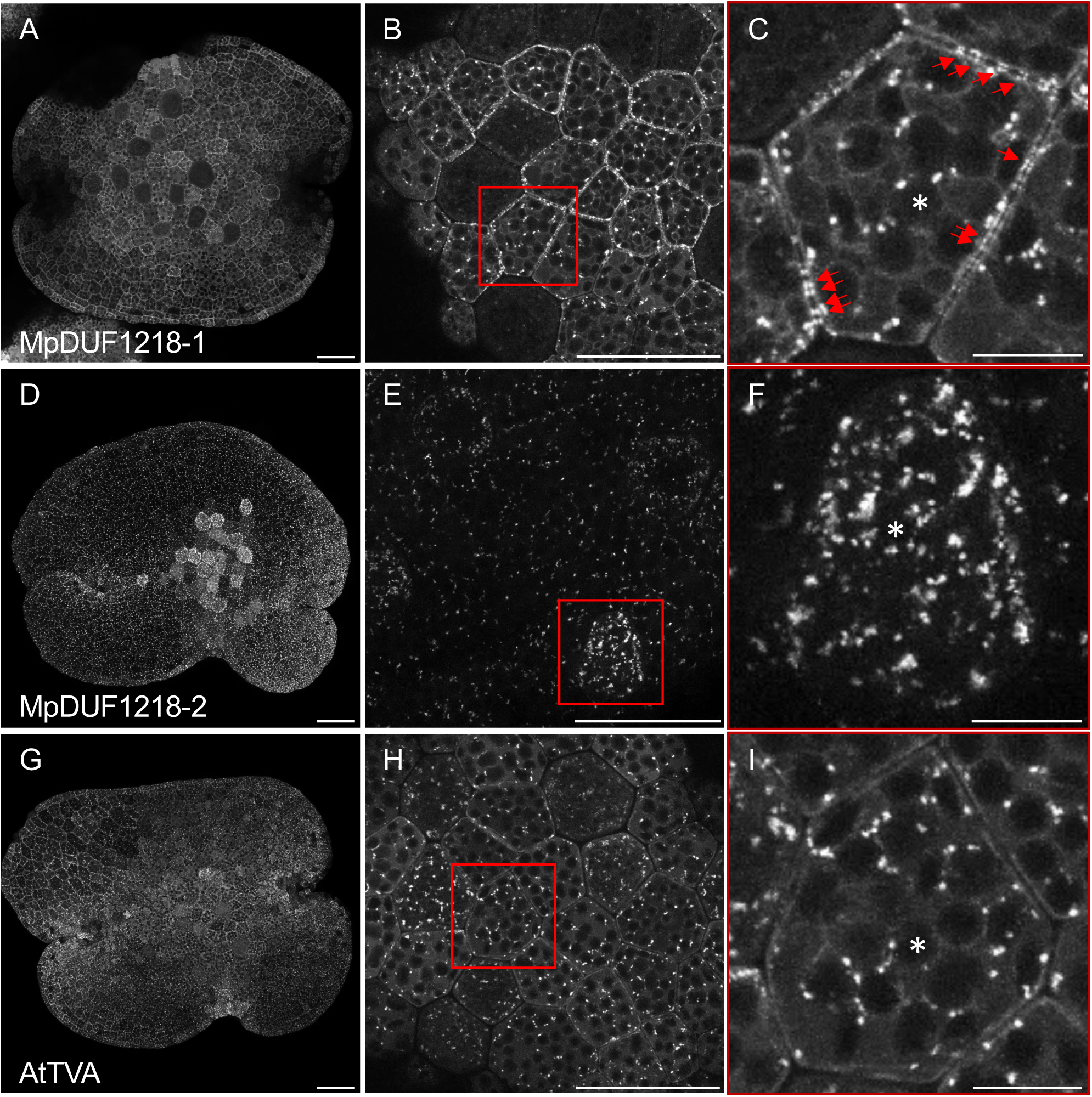
Localization of p*35S*::MpDUF1218-1-, MpDUF1218-2-, amd AtTVA-mCitrine in *Marchantia polymorpha* dormant gemma. (A, D, G) Overview of the entire gemma. (B, E, H) Localization of each protein in the epidermal cells of the dormant gemma. All three proteins form puncta inside cells. (C, F, I) Four-fold enlarged images of the red boxed sections in B, E, and H. Asterisks mark chloroplasts - hollow areas inside cells without fluorescent signals - which are less apparent in MpDUF1218-2, due to its weaker cytoplasmic fluorescent signals. Red arrows point to juxtapose accumulation of proteins on the plasma membrane. Scale bar, 50 μm in (A, B, D, E, G, H); 10 μm in (C, F, I).

To visualize plasmodesmata, we stained the transgenic plants with the aniline blue dye. Because callose staining of dormant gemmae is highly variable and the staining becomes more stable after the development of air pores (Hsu *et al*., 2025), we tested the colocalization of the three DUFs in 11-day-old *M. polymorpha* thalli in which air pores had developed (Fig. 4A-L). As suspected from the fluorescence images in Fig. 3, only MpDUF1218-1-mCitrine consistently colocalized with the aniline blue fluorescence, indicating its plasmodesmata association (Fig. 3A-D, red arrows). This observation was confirmed by quantifing the signals by measuring and plotting the fluorescent intensity of both the green and blue channels, according to their position along the cell wall (Fig. 3M). We interpret that in *N. benthamiana* and *M. polymorpha*, MpDUF1218-1 exhibits plasmodesmata localization, indicating the presence of a shared plasmodesmata-targeting mechanism in both species. The plasmodesmata localization of AtTVA in *N*.

**Figure 4.**
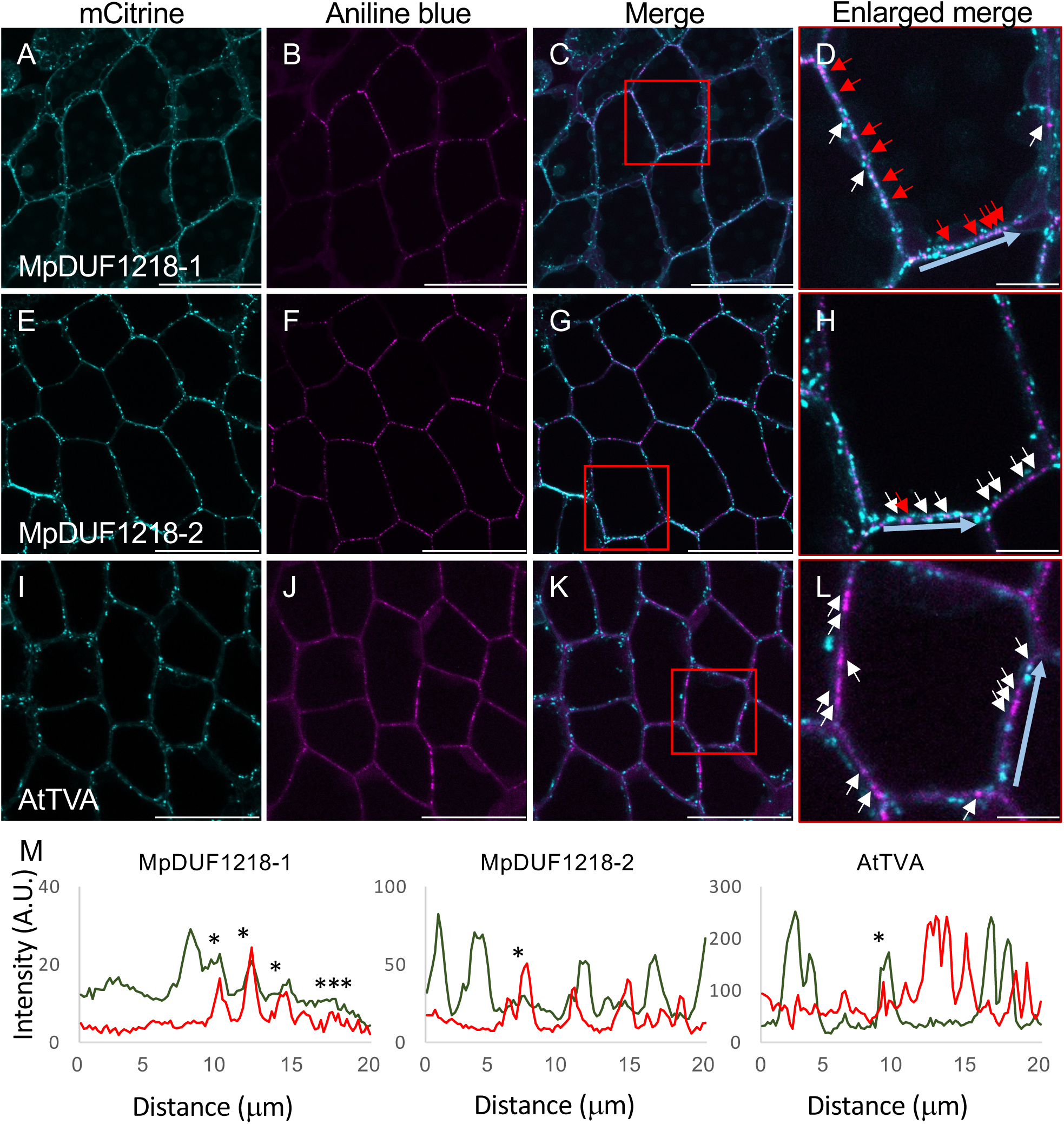
Colocalization of p*35S*::MpDUF1218-1-, MpDUF1218-2-, and AtTVA-mCitrine with aniline blue staining in *Marchantia polymorpha* 11-day-old thalli. (A, E, I) Fluorescent signals of the indicated fusion proteins expressed in the epidermal cells of *M. polymorpha* 11-day-old thalli. (B, F, J) Aniline blue staining was used as the plasmodesmata marker. (C, G, K) The merged images of the previous two panels are shown respectively. (D, H, L) Three-fold enlarged images of the red boxed areas in C, G, and K. Red arrows point to colocalization of the candidate proteins with aniline blue staining. White arrows point to aniline blue signals only. Scale bar, 50 μm for all panels except D, H, L (10 μm). (M) Intensity plot of the mCitrine fusion proteins (green lines) and aniline blue (red lines) measured along the cell wall indicated by the blue arrow in D, H, and L. The x-axis is the distance along the cell wall in μm; the y-axis is the mean intensity of blue and red channels in arbitrary units (A. U.). Asterisks indicate the positions of overlapping peaks.

### Phylogenetic analysis of the DUF1218 domain proteins indicates a divergence occurred after the emergence of Gymnosperms

The targeting of AtTVA to plasmodesmata in a species-specific manner suggests that DUF1218 domain proteins underwent evolutionary adaptation. To understand this process, we performed a deep phylogenetic analysis (Fig. S3) using previously established methods (Mutte *et al*., 2018, Mutte and Weijers, 2020). We found that DUF1218 domain proteins originated from Charophyte algae and could be categorized into three major clades - the AtTVA-, MpDUF1218-3- and MpDUF1218-4-containing clades (Fig. S2). The AtTVA-clade DUF1218 domain proteins (Fig. S3, purple branches) remained as single copies in the ancestors of Bryophytes, Lycophytes and Ferns. It diverged into three clades in seed plants. A summary illustration is presented in Fig. 5A. It is worth noting that the hornworts (represented by *Anthoceros agrestis*) lost the DUF1218 domain protein (Fig 5A). Reciprocal BLAST results of all bryophytes and lycophyte homologs confirmed homology with AtTVA or At4G27435, indicating that the AtTVA-clade represents the ancestral clade of this protein family (Fig. S3). The two copies of MpDUF1218 proteins, MpDUF1218-1 and MpDUF1218-2, are a result of duplication within *M. polymorpha* and belong to the same clade. The duplications in dicots and liverworts might represent a divergence that affects the plasmodesmata localization. Furthermore, MpDUF1218-1 has the highest homology to AtTVA, while MpDUF1218-2 is more similar to another AtDUF1218 protein (At4G27435). This may suggest a common feature required for plasmodesmata localization in AtTVA and MpDUF1218-1 is absent in MpDUF1218-2. Our hypothesis, that AtTVA might have obtained a different plasmodesmata-targeting element, which evolved later and can only be recognized in vascular plants, is supported by the divergence.

**Figure 5.**
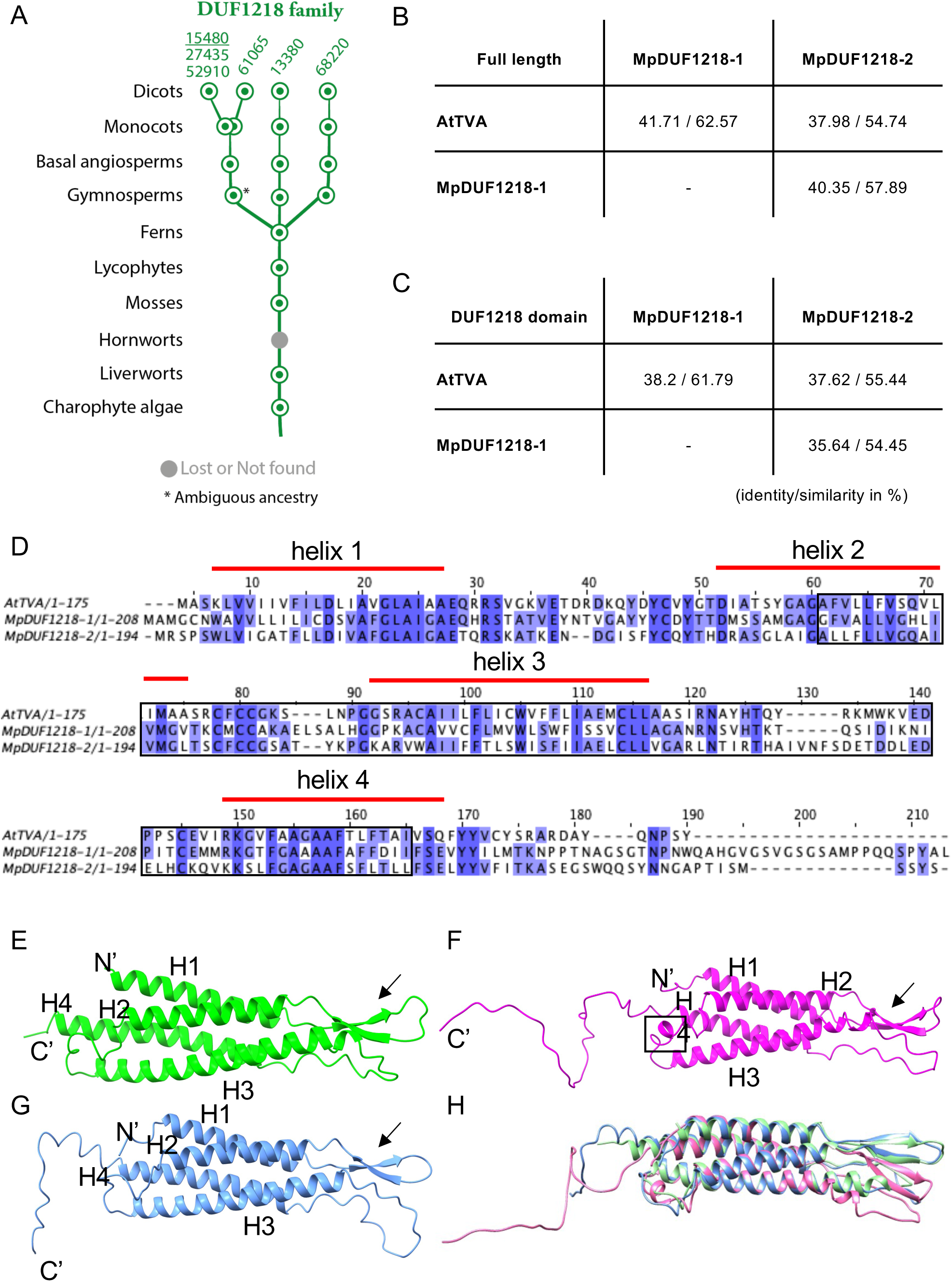
DUF1218 domain proteins diverged after the evolution of Gymnosperms. (A) A summarized evolution road map of DUF1218 domain proteins. All DUF1218 proteins remain as single copy in the ancestral states in Bryophytes, Lycophytes and Ferns and diverge into different branches in seed plants. The numbers on top of the branches indicate the gene ID number of each Arabidopsis DUF1218 protein; the ID number of AtTVA is underlined. (B) Identity and homology of full-length AtTVA, MpDUF1218-1 and MpDUF1218-2 and (C) DUF1218 domains are shown as percentage values. (D) Amino acid sequence alignment of AtTVA, MpDUF1218-1, and MpDUF1218-2. Dark blue boxes indicate identical amino acids, and light blue boxes indicate amino acids with similar properties. Dashes indicate no matching amino acids in the position. Red bars above the sequences show the position of each helix. Black box marks the region of predicted DUF1218 domain, analyzed by HMMER (https://www.ebi.ac.uk/Tools/hmmer/). (E-G) Predicted protein structures of (E) AtTVA, (F) MpDUF1218-1, and (G) MpDUF1218-2. (H) The superimposed images of all three proteins. N’, N-terminal; C’, C-terminal, H1-H4, the four predicted a-helix structures. Black arrows point to the b-sheet structure.

### Amino acid sequence rather than protein structure might be critical for plasmodesmata localization

To explore the potential plasmodesmata-targeting elements in MpDUF1218-1 and AtTVA, we compared the amino acid sequence of AtTVA, MpDUF1218-1, and MpDUF1218-2 (Fig. 5B-D). The full-length AtTVA protein had 41.71% and 37.98% identity and 62.57% and 54.74% similarity to MpDUF1218-1 and MpDUF1218-2, respectively. When we focused on the DUF1218 domain, AtTVA identity to MpDUF1218-1 and MpDUF1218-2 was 38.2% and 37.62%, while similarity was 61.79% and 55.44%, respectively. Similarly, between the two MpDUF1218 proteins, the full-length sequence shared 40.35% identity and 57.89% similarity, while identity and similarity between the DUF1218 domains were 35.64% and 54.45%, respectively (Fig. 5B-D).

The variations in amino acid sequences may affect the overall structure of a protein and consequently its localization and function. We predicted the structures of all three proteins by the Alphafold server (https://alphafoldserver.com/). 4 α-helices and a short antiparallel β-sheets between helix 1 and helix 2 are present in all three protein predictions (Fig 5E-H). An additional small helical structure is present between helix 2 and 3 in MpDUF1218-1 (Fig. 5F, black square). Superimposition of the three proteins by ChimeraX program (https://www.cgl.ucsf.edu/chimerax/; Pettersen *et al*., 2021) reveals that beside the variable lengths of the C-terminal region and the additional small helix in MpDUF1218-1, all three proteins have high structural similarity (Fig. 5H). We calculated the Root Mean Square Deviation (RMSD) of atom pairs between all three proteins by the ChimeraX. When the RMSD is smaller than two angstroms (Å), the two structures are considered highly similar (Jewett *et al*., 2003, Armougom *et al*., 2006). The RMSD of 107 pruned atom pairs between AtTVA and MpDUF1218-1 was 0.861Å, of 139 pruned atom pairs between AtTVA and MpDUF1218-2 was 0.847 Å, and of 108 pruned atom pairs was 0.870 Å. The RMSD between all three proteins were below 2 Å, suggesting that the sequence variations did not profoundly alter the structure. Therefore, plasmodesmata-targeting elements might be specific sequences instead of overall structural variations.

### Helix 1 and Helix 2 of MpDUF1218-1 carry plasmodesmata-targeting information in

Since the plasmodesmata-targeting elements might be specific sequences, we performed a helix-swap (HS) analysis between MpDUF1218-1 and -2 to identify the crucial elements responsible for plasmodesmata localization in MpDUF1218-1. We generated four chimeric MpDUF1218-1 constructs in which each helix (helix 1 to helix 4 in Fig. 6A) was substituted by the helix of MpDUF1218-2 at the corresponding position, indicated as MpDUF1218-1 HS1, HS2, HS3, and HS4, respectively. We found that the plasmodesmata localization of the proteins was lost in both MpDUF1218-1 HS1 and HS2 transgenic plants (Fig. 6B, C). Furthermore, in MpDUF1218-1 HS1-mCitrine the formation of puncta was largely reduced, while in MpDUF1218-1 HS2-mCitrine, the puncta were still clearly observed (Fig. 6B, C). MpDUF1218-1 HS3 and MpDUF1218-1 HS4 did not interfere with plasmodesmata localization of the chimeric MpDUF1218-1 proteins (Fig. 6D, E, red arrows). Our results revealed that a key regulatory feature for plasmodesmata localization in *M. polymorpha* is positioned within the first two helix regions of MpDUF1218-1.

**Figure 6.**
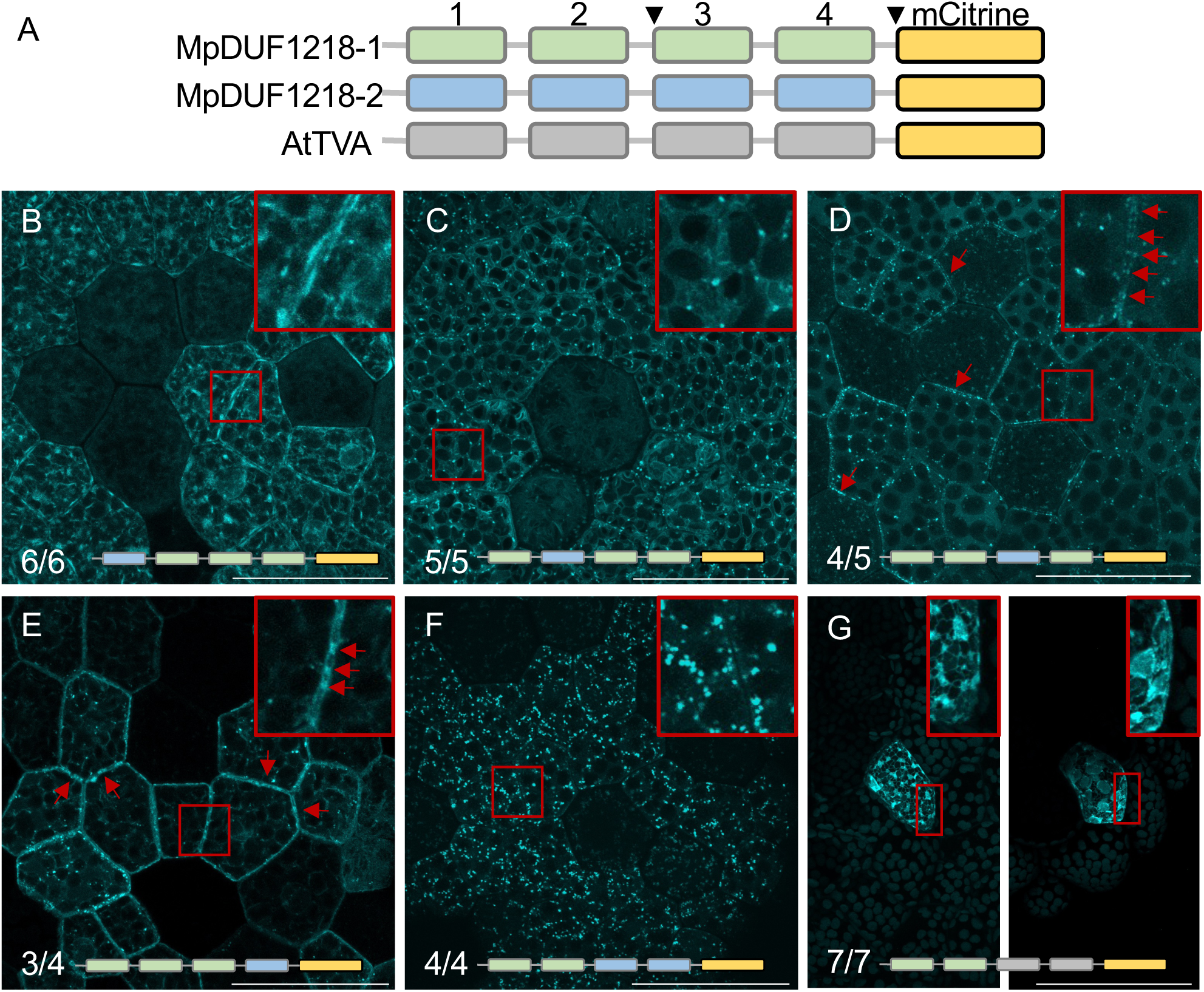
The first two helices are critical for the plasmodesmata localization of MpDUF1218-1-mCtrine in *Marchantia polymorpha*. (A) Cartoon of the three DUF1218 proteins. Blocks 1-4 represent helices with the respective number above. Visual marker mCitrine (yellow box) was added to the C-terminus of all three proteins. When consecutive helices 3 and 4 were simultaneously swapped, the entire sequence between the arrowheads was swapped. (B-G) Fluorescent signals of the indicated helix-swapped (HS) fusion proteins as indicated at the bottom of each panel. Three-fold enlarged images of the red boxed sections are shown in the upper right corner of each panel. (B-F) Helices were swapped between MpDUF1218-1 (green) and MpDUF1218-2 (blue) (B) MpDUF1218-1 HS1 signals were observed mostly inside cells with string-like enrichments at cell peripheries. (C) MpDUF1218-1 HS2 formed puncta inside cells. (D, E) MpDUF1218-1 HS3 and MpDUF1218-1 HS4 formed puncta inside cells with juxtaposition puncta at cell peripheries (red arrows). (F) Constructs carry 2 swapped helices from MpDUF1218-1 HS3,4 (helix-swapped 3 and 4), forming cytosolic puncta structures without clear juxtaposition puncta. (G) Helices 3 and 4 were swapped between MpDUF1218-1 (green) and AtTVA (grey). Clear network structures were formed, representing the ER structure without obvious puncta signals at cell peripheries. Scale bar, 50 μm.

We further tested if helix 1 and helix 2 of MpDUF1218-1 are sufficient to guide the plasmodesmata localization. We swapped both helix 3 and helix 4 of MpDUF1218-1 with the corresponding helices of MpDUF1218-2 (MpDUF1218-1 HS3,4; the entire sequence between helix 3 and the end of the protein was swapped, indicated by arrowheads in Fig. 6A). In the transgenic *M. polymorpha*, we noticed clear puncta structures localized mostly at the cytoplasm, resembling the puncta of the original MpDUF1218-2 (Fig. 6F and 3H). To confirm whether these puncta can localize to plasmodesmata, we performed the aniline blue staining at 11-day-old thalli and found that about 8% of the MpDUF1218-1 HS3,4-mCitrine particles overlapped with aniline blue signals (Fig. S4). These data indicate that the first two helices of MpDUF1218-1 are essential but not sufficient for plasmodesmata localization. Alternatively, helix 3 and helix 4 of MpDUF1218-2 may have a stronger puncta-formation property than those of MpDUF1218-1, causing the protein to form intracellular puncta that may mask or override the plasmodesmata targeting.

To test the alternative possibilities, we constructed a similar helix 3 and 4 swap between MpDUF1218-1 and AtTVA. We reasoned that the AtTVA-mCitrine signals were also observed at the plasma membrane in the transgenic plants (Fig. 3A-C); therefore, helix 3 and 4 of AtTVA might have less puncta formation tendency compared to those of MpDUF1218-2. By using an agrobacterium-based transient expression assay, we successfully detected a few cells expressing the MpDUF1218-1 HS3,4-mCitrine protein. Unexpectedly, we observed clear network-like structures resembling the cortical ER (Fig. 6G). This result indicated that, even though helix 1 and helix 2 are important for MpDUF1218-1 to target plasmodesmata, the rest of the protein also influenced the sorting process.

### Helix 1 of MpDUF1218-1 is also important for plasmodesmata targeting in *N. benthamiana*

To investigate whether the plasmodesmata-targeting mechanism for MpDUF1218-1 is conserved in vascular plants, we expressed the helix-swapped constructs together with the plasmodesmata marker PDLP1-mCherry by agro-infiltration in *N. benthamiana* (Fig. 7). Similar to the observations in *M. polymorpha*, plasmodesmata localization of MpDUF1218-1 HS1-mCitrine was largely reduced (Fig. 7A-D, white arrows). We also observed MpDUF1218-1 HS2-mCitrine with reduced plasmodesmata localization, however, the reduction was not as strong as HS1 (Fig. 7E-H). Furthermore, as in *M. polymorpha*, MpDUF1218-1 HS3 and HS4-mCitrine remained to be localized to plasmodesmata (Fig. 6I-P, red arrows). To clearly compare the alternation of plasmodesmata localization, we established a colocalization analysis method to compare the changes between all domain-swapped proteins (Fig 7Q). We use the ubiquitously localized mCitrine protein as a negative control and the plasmodesmata labeling aniline blue staining as the positive control to perform the colocalization analysis with PDLP1-mCherry. We obtained 13.5±8.3% colocalization between mCitrine and PDLP1-mCherry and 82.9±2.1% colocalization between aniline blue and PDLP1-mCherry. For MuDUF1218-1 series, the full length was 52.6±3.6% colocalized with PDLP1-mCherry and all domain swap affected the colocalization percentage. HS1 significantly reduced the colocalization (18.5±2.3%), HS2 (40.26±16.22%) and HS4 (37.6±5.4%) also caused the reduction of colocalization percentage but the reductions were not statistically significant. Similarly, HS3 (62.8±3.5%) increased the colocalization with PDLP1-mCherry but it was not statistically significant (Fig. 7Q, Supplemental Data S1).

**Figure 7.**
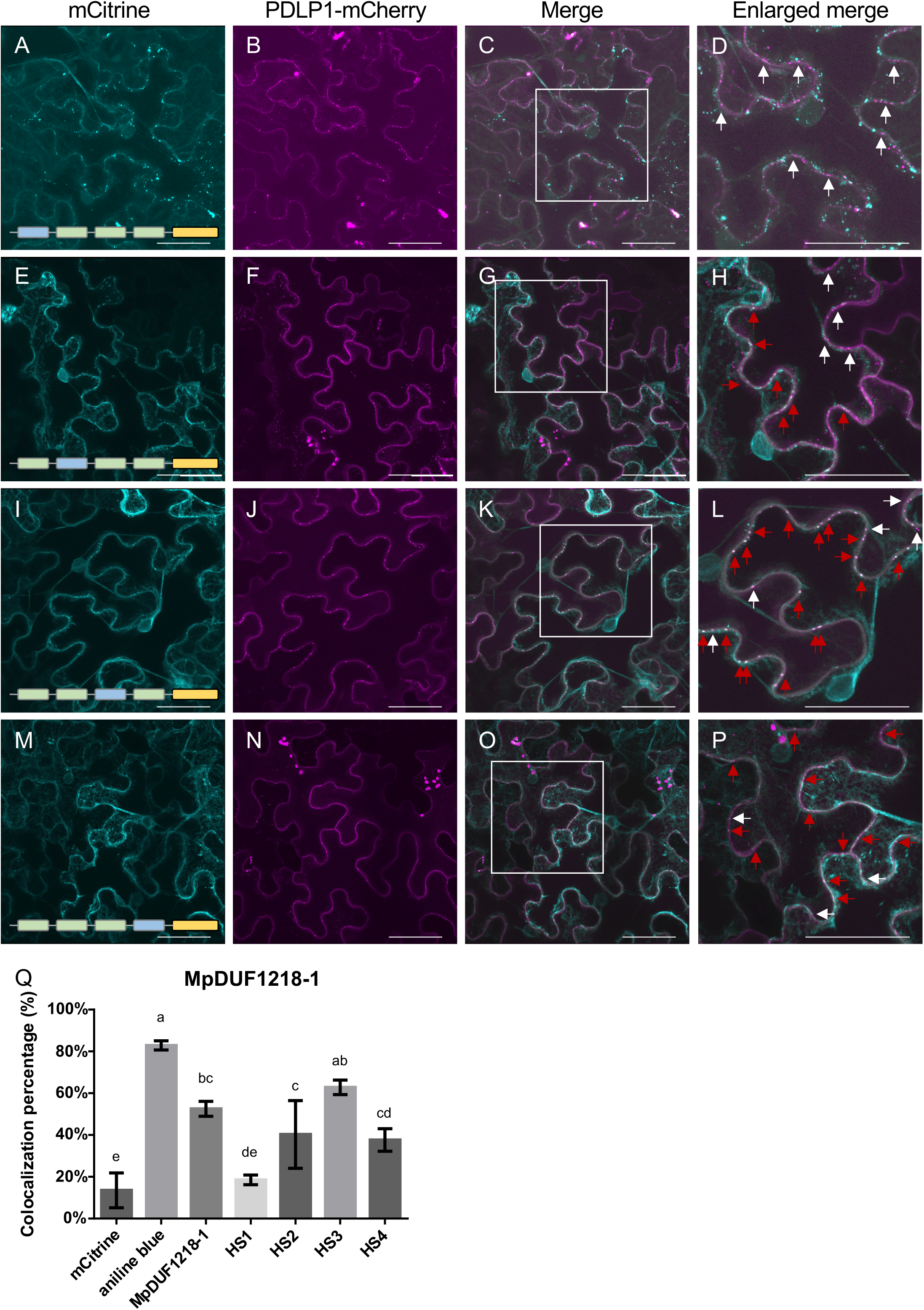
The first two helices are also critical for the plasmodesmata localization of MpDUF1218-1-mCitrine in Nicotiana benthamiana. (A, E, I, M) Fluorescent signals are presented from the indicated helix-swapped proteins at the bottom of each panel expressed in the leaf epidermal cells of N. benthamiana. The composition of the tested proteins is shown in Fig. 6A. Swapping involved helices from MpDUF1218-2 (blue) into the test construct MpDUF1218-1 (green) (B, F, J, N) Plasmodesmata were visualized using p35S::PDLP1-mCherry expressed on the same leaf sections. (C, G, K, O) Merged images of the previous two panels, respectively. (D, H, L, P) Two-fold enlarged images from the red boxed sections in C, G, K, and O. Red arrows point to the colocalization of the helix-swapped proteins with the plasmodesmata marker. White arrows point to the plasmodesmata markers only. Note that in the merged images, the plasmodesmata marker alone showed magenta color (C, D, G, H, S, T), while colocalized with the helix-swapped proteins and the plasmodesmata markers showed white color (K, L, O, P). Scale bar, 50 mm. (Q) Colocalization analysis of all MpDUF1218-1 single domain-swap constructs with PDLP1-mCherry. mCitrine was used as the negative control while aniline blue staining was used as the positive control. n=3 for each construct. The data were analyzed by one-way ANOVA with Tukey’s HSD post hoc analysis.

We also expressed the MpDUF1218-1 HS3,4-mCitrine (with MpDUF1218-2) and MpDUF1218-1 HS3,4-mCitrine (with AtTVA) in *N. benthamiana* and observed plasmodesmata localization of the chimeric proteins (Fig. S5). Similar to the subcellular localization in *M. polymorpha*, when combined with MpDUF1218-2 helix 3 and 4, the chimeric formed cytoplasmic puncta with few overlaps with the PDLP1-mCherry signals (Fig. S5B-E and S5J), while the combination with AtTVA helix 3 and 4, the chimeric proteins became more membrane localized with higher colocalization percentage with PDLP1-mCherry (Fig. S5F-I, and S5J).

Additionally, we constructed MpDUF1218-1 H1 and MpDUF1218-1 H2-mCitrine fusion proteins and expressed them in *N. benthamiana* to test whether these helices individually could target plasmodesmata. Both fusion proteins did not localize at plasmodesmata as they obtained similar PDLP1-mCherry colocalization percentage with the negative control (H1: 17.3±2%; H2: 18.2±3.2%, Fig. S6A-H and S6M, Supplemental Data S1). We further constructed a MpDUF1218-1 H1 and H2-mCitrine (MpDUF1218-1 H1,2-mCitrine) fusion protein to observe its subcellular localization. The MpDUF1218-1 H1,2-mCitrine proteins were mostly localized at ER, with some fluorescent signals colocalized with the PDLP1-mCherry signals (36.3±18.2%, Fig. S6I-M, Supplemental Data S1). These data suggest that MpDUF1218-1 H1 and H2 collectively contain plasmodesmata-targeting elements and can be recognized by the plasmodesmata-targeting mechanism in *N. benthamiana*. The comparable localizations of the chimeric proteins between *M. polymorpha* and *N. benthamiana* support the idea that MpDUF1218-1 may utilize the conserved plasmodesmata-targeting mechanism to locate to plasmodesmata in both species.

### Helix 2 and helix 3 deletion disrupted the plasmodesmata localization of AtTVA

As AtTVA localizes at plasmodesmata only in *N. benthamiana*, this plant could possess another plasmodesmata-targeting mechanism different from the one that recognizes MpDUF1218-1. To explore this possibility, we analyzed helices that could be crucial for the plasmodesmata localization of AtTVA. Helix deletion was our approach here to avoid potential interference with protein domains from other species. We transiently expressed the AtTVAΔH1-, ΔH2-, ΔH3-, and ΔH4-mCitrine in *N. benthamiana* by agro-infiltration (Fig. 8). We observed colocalizations with the plasmodesmata marker, PDLP1-mCherry, indicating AtTVAΔH1- and AtTVAΔH4-mCitrine were targeted to plasmodesmata (Fig. 7A-D and M-P). However, AtTVAΔH2- and AtTVAΔH3-mCitrine had limited colocalization with PDLP1-mCherry at plasmodesmata (Fig. 7E-L). To evaluate the colocalization with PDLP1-mCherry, we also performed the colocalization per centage analysis as previously described. The data were consistent with the observation that AtTVAΔH1-(49.1±5.7%) and AtTVAΔH4-mCitrine (43.2±5.3%) showed similar colocalization percentage with PDLP1-mCherry to the full-length AtTVA (42.1±3.8%). AtTVAΔH2-(25.6±4.6%) and AtTVAΔH3-mCitrine (23.6±3.3%) had limited colocalization with PDLP1-mCherry (Supplemental Data S1). Our data showed that the plasmodesmata-targeting element maps to helix 2 and 3 of AtTVA, which is distinct from that of MpDUF1218-1, supporting the idea that a separate plasmodesmata-targeting mechanism exists in *N. benthamiana* but is absent in *M. polymorph*a.

**Figure 8.**
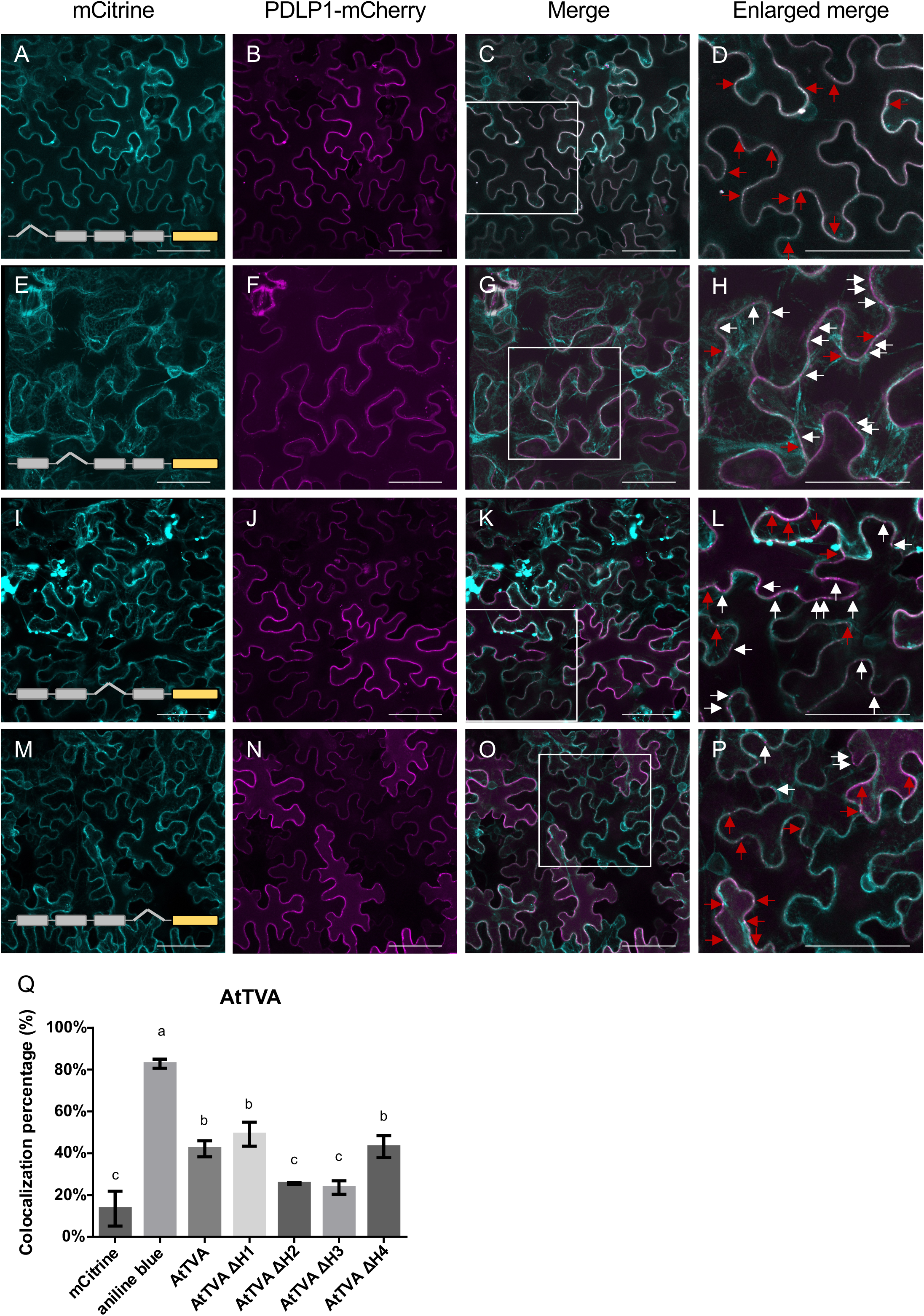
Deletion of AtTVA Helix 2 and 3 resulted in the loss of its plasmodesmata localization. (A, E, I, M) Fluorescent signals of the indicated fusion proteins expressed in the leaf epidermal cells of *N. benthamiana*. The composition of the tested proteins is shown in Fig. 6A. The roof triangle indicates deletion. (B, F, J, N) Plasmodesmata were visualized using p*35S*::PDLP1-mCherry expressed on the same leaf sections. (C, G, K, O) Merged images of the previous two panels, respectively. (D, H, L, P) Two-fold enlarged images of the red boxed sections in C, G, K, and O. In the merged images, plasmodesmata markers alone show magenta color, when colocalized with the helix-deleted proteins, the plasmodesmata markers showed white color (C, D, G, H, K, L, O, P). Red arrows point to the colocalization of the candidate proteins with the plasmodesmata markers. White arrows point to the plasmodesmata markers only. Scale bar, 50 mm. (Q) Colocalization analysis of AtTVA domain-deletion constructs with PDLP1-mCherry. mCitrine was used as the negative control while aniline blue staining was used as the positive control. Note that the same mCitrine and aniline blue data were reused as a comparison reference. n=3 for each construct. The data were analyzed by one-way ANOVA with Tukey’s HSD post hoc analysis.

## Discussion

### At least two plasmodesmata-targeting mechanisms exist in *N. benthamiana*

Plasmodesmata are key intercellular communication channels that evolved after plants inhabited land. We hypothesize that while plasmodesmata are found in all land plants, different strategies to target their proteins to the structure may have evolved. We investigated three plasmodesmata-targeting proteins from two different plants and tested them in two species (*N. benthamiana* and *M. polymorpha),* where we obtained evidence in support of our hypothesis. AtTVA and its homolog MpDUF1218-1 are plasmodesma-targeted in *N. benthamiana*; however, only MpDUF1218-1 but not AtTVA localized to plasmodesmata in *M. polymorpha*. The consistent plasmodesmata localization of MpDUF1218-1 supports the idea that a conserved plasmodesmata-targeting system is present in both *N. benthamiana* and *M. polymorpha*. The distinct localization of AtTVA indicates that the plasmodesmata-targeting system for AtTVA in *N. benthamiana* is absent in *M. polymorpha*. Our phylogenetic analysis indicated that DUF1218 domain proteins of AtTVA clade diverged after the emergence of gymnosperm. It is possible that DUF1218 domain proteins in gymnosperm and angiosperm might have been reshaped to adopt a new mechanism. The similar structure of all three proteins reduced the possibility that the evolution event caused a structural alternation and allow a different plasmodesmata-targeting mechanism to recognize the DUF1218 domain proteins in gymnosperm and angiosperm. The dissection of cis-elements showed that the first two helices of MpDUF1218-1 are critical for its plasmodesmata localization. On the contrary, the plasmodesmata localization of AtTVA requires the second and third helixes. The different plasmodesmata-targeting elements, which are present in MpDUF1218-1 and AtTVA make the existence of different plasmodesmata-targeting systems in *N. benthamiana* and *M. polymorpha* very likely.

Interaction with specific lipid rafts located in plasmodesmata could be one of the potential regulations for plasmodesmata targeting. It has been shown that the sterols and complex sphingolipids are enriched in the plasmodesmata-plasma membrane domain (Grison *et al*., 2015). A few plasmodesmata localized proteins, like Multiple C2 domain and Transmembrane region proteins (MCTPs) and Synaptotagmins (SYTA), contain lipid binding domain and have been suggested to localize to plasmodesmata by specifically interacting with the lipids in plasmodesmata (Vaddepalli *et al*., 2014, Yuan *et al*., 2018, Brault *et al*., 2019). In a recent study, the second C2 domain of MCTP4 has been demonstrated to specifically interact with phosphatidy-linositol-4-phosphate (PI4P) (Perez-Sancho *et al*., 2025). The interaction is also crucial for the plasmodesmata localization of MCTP proteins. The inhibition of PI4P synthesis by phenylarsine oxide (PAO, a PI4 kinase inhibitor) treatment reduced the accumulation of MCTP4 at plasmodesmata, while in the s*uppressor of actin* 7 (*sac7*, a PI4P phosphatase) mutant, the increase of PI4P in plasmodesmata enhanced the accumulation of MCTP4 (Brault *et al*., 2019). Furthermore, plasmodesmata callose binding protein 1 (PDCB1) and plasmodesmata localized β-1,3-glucanase 2 (PdBG2) contain GPI-anchoring domains, their localizations were altered under the treatment of sterol inhibitors (Grison *et al*., 2015).

As plasmodesmata-targeting elements in MpDUF1218-1 and AtTVA are predicted as transmembrane α-helixes, they might perform different lipid interactions with the aforementioned lipid-binding domain proteins. Different helices of DUF1218 might have different hydrophobicity and can interact with different lipids, causing diverse subcellular localizations. Previously, PLASMODESMATA LOCLIZATION PROTEIN 1 (PDLP1) has been demonstrated to utilize its transmembrane domain for plasmodesmata targeting. When the C-terminal transmembrane domain fused with a Citrine fluorescent protein, the fusion protein was localized at plasmodesmata (Thomas *et al*., 2008). Mutation of the amino acid valine to alanine (V288A) located towards the C-terminal part of the transmembrane domain can alter the localization of the fusion protein, suggesting that the hydrophobicity of the transmembrane domain is crucial for plasmodesmata targeting (Thomas *et al*., 2008). Because different lipid bilayers are composed of fatty acids with aliphatic chains of varying lengths, the length of the transmembrane domain or the overall protein structure may influence the ability to locate to certain lipid domains (Nyholm, 2015). Within the DUF1218 proteins, the predicted helix domains as well as the predicted protein structures resemble each other (Fig 4C-H); the number of aliphatic amino acids in each helix of MpDUF1218-1 and -2 is very similar, but the subcellular localization is distinct. One possible explanation is that a sequence-based sorting system might be responsible, a hypothesis that needs experimental verification.

In addition, several inter-helix sequences of MpDUF1218-1 were not tested for their roles in plasmodesmata targeting. In the helix-swap analysis, we retained all the inter-helix sequences from MpDUF1218-1 but observed that only MpDUF1218-1 HS1 and MpDUF1218-1 HS2 lost plasmodesmata localization. This result suggests that disruption of the sequence following helix 2 does not affect the localization of MpDUF1218-1 to plasmodesmata (Fig. 6 and 7). Additionally, the single MpDUF1218-1 H1 and MpDUF1218-1 H2 failed to localize at plasmodesmata while the MpDUF1218-1 H1H2 localized at plasmodesmata suggests that the linker between helix 1 and helix 2 might also contributes to MpDUF1218-1’s targeting.

### Subcellular localization of DUF1218 proteins is determined by different helices

Besides targeting to plasmodesmata, we also observed a variety of subcellular localizations of DUF1218 proteins, including plasma membrane, ER, as well as forming puncta structures. For multiple transmembrane domain proteins, the sorting mechanism is still vague. In our study, the transient expression MpDUF1218-1 H1H2-mCitrine construct localized primarily at ER with limited puncta (Fig. S6I), indicating that the presence of both helixes provides strong ER retention properties. Furthermore, the full-length MpDUF1218-1 formed few puncta but had plasma membrane and plasmodesmata localization, suggesting that with helix 3 and helix 4, the MpDUF1218-1 protein is translocated from ER to plasma membrane, maybe via the puncta. Once the protein locates on the plasma membrane, the first two helixes will provide the information for the protein to target plasmodesmata.

Different subcellular localization properties seem to be present in different DUF1218 proteins, as the combination of MpDUF1218-1 HS3,4 (swapped with MpDUF1218-2; Fig S5B, J) formed strong puncta structures but few plasmodesmata accumulation, indicating that MpDUF1218-2 helix 3 and 4 may have a dominant condensation function. It is worth noting that the subcellular localizations of most chimeric MpDUF1218s are comparable between *M. polymorpha* and *N. benthamiana*, suggesting the conservation of the sorting mechanisms. Unexpectedly, the chimeric MpDUF1218-1 HS3, 4 (swapped with AtTVA) localized at both ER and plasmodesmata in *N. benthamiana*, but showed strong ER retention properties but no plasmodesmata localization in *M. polymorpha* (Fig. 6G), in contrast to the partial plasmodesmata localization status in *N. benthamiana* (Fig S5F, J). This phenomenon again supports the idea that the plasmodesmata localization regulation might be different between two plant species but also shows that the distribution of DUF1218 proteins between puncta, ER or plasma membrane is determined not only by one or two helixes but the sum of the entire protein.

In summary, our study reveals that at least two membrane-based plasmodesmata-targeting systems are present to direct plasmodesmata-related transmembrane proteins. These proteins might use different sequence specific transmembrane domains to provide the specificity of plasmodesmata localization. However, it may be the overall properties of the whole transmembrane domains that is regulating the fine-tuning with directing a proportion of proteins to the ER, some to form puncta in the cytoplasm and others to enriched at plasmodesmata. Further structural prediction, amino acid mutagenesis, and lipid interaction analysis will aid in providing detailed information on the mechanisms by which a plant cell guides these plasmodesmata-targeting proteins to their destinations.

## Materials and Methods

### Plant growth and transformation

*Nicotiana benthamiana* was grown in 10 cm pots in a 1:1 ratio of potting mixture and vermiculite at 25° C under long-day conditions (16 h/8 h light/dark cycles). *Marchantia polymorpha* Takaragaike-1 (Tak-1) was used as the wild type and for generating all transgenic lines. Plants were cultured on ½ B5 plates (1/2 Gamborg’s B5 medium, 0.5 g/L MES, pH 5.7 and 1% agar) in a growth chamber (F-740, HiPoint), at 22°C under a 16h/8h day/night cycles, with 60 μEm−2s−1 LED white light.

Transgenic *M. polymorpha* plants were generated via the agrobacterium-mediated transformation method (Ishizaki et al., 2008; Kubota et al., 2013). In brief, after removing the apical meristem area of 14-day-old Tak-1 thallus, the thallus was cut into small pieces then incubated on ½ B5 medium agar plates with 1% sucrose for 3 days. Agrobacteria the respective target plasmid was cultured in LB medium with corresponding antibiotics for 2 days, harvested by centrifugation, resuspended in 0M51C medium with 100 μM acetosyringone (3,5-dimethoxy-4-hydroxyacetophenone), diluted to OD600=1.0, and cultured on benchtop for 6 hrs. 15 to 20 regenerating plantlets and 1 mL of agrobacteria (OD600= 1.0) were cocultured in 50 mL 0M51C medium with 100 μM acetosyringone in a growth chamber under a 16h/8h day/night cycle with agitation at 100 rpm at 22° C for 3 days. After washing the transformed thalli 5-6 times with sterilized, they were plated on ½ B5 plates containing antibiotics for selection (10 mg/L hygromycin B and 100 mg/L cefotaxime). The successfully transformed plants were designated as T1 plants. G1 gemmae harvested from the T1 generation were cultured on ½ B5 plates containing the antibiotic selection. G2 gemmae, which were harvested from the G1 generation, were used for analyses.

### Plasmid construction

p35S::Mp2g06670-, Mp2g17160-, Mp2g17160-, and MpTET3a-mGFP5 were generated by PCR amplification with primers listed in Supplemental Table S1 and cloned into pGEM-T Easy vector (Promega) for sequence confirmation. Target genes were further subcloned into the pEpyon-32H binary vector obtained from Prof. Chang-Hsien Yang (Graduate Institute of Biotechnology, National Chung Hsing University, Taichung, Taiwan), which contains a mGFP5 fluorescent protein (Dai *et al*., 2019), with the restriction enzymes indicated in the primers’ name.

p35S::AtTVA-, MpDUF1218-1-, MpDUF1218-2-, MpDUF1218-1 H1-, H2-, and H1H2-mCitrine constructs were generated by homology-based SLiCE reaction (Zhang et al., 2014) and Gateway LR reaction (Invitrogen). Gene sequences corresponding to AtTVA, MpDUF1218-1 and MPDUF1218-2 with extra 15 base pairs homologous to the 5’ half and 3’ half LIC sequence were amplified from Arabidopsis Col-0 cDNA and *M. polymorpha* Tak-1 cDNA, using gene-specific primers listed in Supplemental Table S1. The PCR products were cloned into Hpa I (NEB)-digested linear pENTR-LIC plasmid by SLiCE reaction and verified by sequencing. The LIC sequence of pENTR-LIC plasmid was generated by primer synthesis according to a previous study (De Rybel *et al*., 2011) and fused to pENTR^TM^/D-TOPO^TM^ vector by TOPO reaction provided from the manufacturer (Thermo Fisher Scientific). Plasmids containing target fragments were recombined into the binary expression vector, pMpGWB106 (Ishizaki et al., 2015), using the Gateway™ LR clonase™ according to the user’s manual (Thermo Fisher Scientific). All helix-swapped constructs were generated by the SLiCE reaction. The corresponding helix domain from MpDUF1218-2 plus extra 15-base pairs homologous to the upstream and downstream sequence of the replaced helix were amplified by PCR with primers listed in Supplemental Table S1. The pENTR-MpDUF1218-1 plasmid was amplified using the primers listed in Supplemental Table S1 to generate the linearized plasmid without the corresponding helix. The two fragments were fused by the SLiCE reaction and the successful entry plasmids were recombined into pMpGWB106 by the Gateway^TM^ LR clonase^TM^.

The AtTVA domain-deletion constructs were generated by PCR the pENTR-AtTVA plasmid with primers listed in Supplemental Table S1. The linear PCR products were ligated by T4 DNA ligase, transformed to *E. coli* and analyzed by colony PCR. The plasmids with corresponding deletion were sequenced and recombined to pMpGWB106 by the Gateway™ LR clonase™.

### Phylogenetic analysis

To identify and extract the homologous proteins of DUF1218 family, representative family members from *Arabidopsis thaliana*, *Marchantia polymorpha*, *Physcomitrium patens*, *Amborella t*richopoda, Ceratopteris richardii, *Zea mays*, *Solanum lycopersicum* and *Oryza sativa* were used to make an HMM profile using HMMER v3 (Finn *et al*., 2011). This profile was used to search for more than 100 proteomes of reference and representative species, as per the methods mentioned earlier (Mutte *et al*., 2018, Mutte and Weijers, 2020). Briefly, the proteomes of the selected species were searched for homologs using HMMER. All the homologs were searched for protein domains using InterProScan (Jones *et al*., 2014). In total, 1313 proteins with the DUF1218 domain (Interpro ID: IPR009606) were selected for further phylogenetic analysis. Sequences were aligned using MAFFT e-ins-i algorithm (v7;Katoh and Standley, 2013) with 1000 iterations followed by trimming with trimAl (Capella-Gutierrez *et al*., 2009) to remove the positions with 0.2 gap threshold. IQ-TREE v2.3.5 (Minh *et al*., 2020) was used to make the phylogeny with a maximum of 100 rapid bootstraps with JTT+R6 as the evolutionary model. Phylogenetic trees were visualized using iTOL server (https://itol.embl.de/).

### Agro-infiltration

Transient expression of all constructs in *N. benthamiana* was performed via the agrobacterium infiltration method (Norkunas et al., 2018). In brief, the agrobacterium GV3101 strain harboring target plasmids was cultured in LB medium with corresponding antibiotics at 28° C for 48 hrs. The harvested bacterium was resuspended in MMA (5 g/L MS salt, 20 g/L sucrose, 2 g/L MES, pH 5.6) medium with 100 μm acetosyringone to OD600=0.5 and incubated at room temperature for 2 hrs. The third to fifth leaves from the top of a 2-week-old plant were infiltrated by placing a 1.0 ml plastic syringe against the abaxial side of the leaves with a finger on the adaxial side for support, applying gentle syringe pressure to infiltrate agrobacteria into the leaves. The infiltrated plants were cultured at 25° C in a growth chamber with 16/8 day/night cycle for 48 hrs and the leaves were subjected to confocal microscopy for observation.

Transient expression of p*35S*::MpDUF1218-1 H3,4-mCitrine (helix-swapped with AtTVA) was performed by the agrobacterium penetration method. Agrobacteria harvesting the target construct was cultured in LB medium with corresponding antibiotics at 28° C for 48 hrs. The agrobacteria were harvested and resuspended in 0M51C medium with 100 μM acetosyringone in a flask, diluted to 50 ml OD600=1.0, and incubated 6 hrs at room temperature on bench. 14-day-old *M. polymorpha* thalli were submerged into the agrobacteria medium and subjected to vacuum for 5 minutes. The medium was cultured with 120 rpm agitation in a growth chamber under long-day conditions (16 h/8 h day/night) at 22° C for 3 days. The thalli were harvested, washed with sterile water for 5 times and cultured on a 1/2 B5 medium plate for another 3 days before confocal microscopy.

### Microscopy

For the observation of protein expression in *N. benthamiana*, a small leaf disc was cut by a paper cutter from an infiltrated leaf and transferred onto a 24 mm x 50 mm coverslip, sandwiched by another 24 mm x 24 mm coverslip and observed under the laser scanning confocal microscope FV3000 (Olympus). mCitrine-containing images were taken with 514 nm excitation and 520-570 nm detection. mCherry-containing images were taken with 561 nm excitation and 600-670 nm detection. For observation of protein expression in *M. polymorpha*, gemmae were picked from the gemmae cup and directly transferred onto a 24 mm x 50 mm coverslip sandwich, sandwiched by another 24 mm x 24 mm coverslip and observed under the laser scanning confocal microscope FV3000 (Olympus). mCitrine-containing images were taken with 514 nm excitation and 520-570 nm detection. For aniline blue staining, images were taken with 405 nm excitation and with 430-470 nm detection.

### Aniline blue staining

For staining N. benthamiana with aniline blue, a small leaf disc was cut by a paper cutter from an infiltrated leaf and transferred into a 1.5 mL eppendorf filled with 200 μL 0.01 mg/ml aniline blue fluorophore (Biosupplise) solution and vacuum (100 mbar) for 5 min in a tabletop vacuum chamber. After vacuum, the leaf disc was further stained for another 30 min before imaging and then transferred onto a slide, observed under the laser scanning confocal microscope FV3000 (Olympus).

For staining *M. polymorpha* with aniline blue, a small piece was cut from 11-day-old thallus and transferred onto a concave cavity slide (Marienfeld) with a drop of 10 μl 0.1 mg/ml aniline blue fluorophore (Biosupplise) solution applied directly on the surface of the thallus. The specimen was incubated for 10 minutes, covered by a cover slide and observed under the laser scanning confocal microscope FV3000 (Olympus).

### Image quantification

All images were analyzed by Fiji (ImageJ, https://imagej.net/software/fiji/).

For Plot Graph, the Plot Profile Tool was used. The area chosen for the colocalization profile was indicated in the figure. The *X*-axis displays the distance from the beginning point to the ending point. The *Y*-axis indicated the average gray value within the area selected.

For colocalization percentage analysis, first, we measured the intensity of cyan and magenta channels in sections along all cell walls in each colocalization analysis image and exported the data into one spreadsheet. We then calculate the intensity average of the two channels independently and select signals that was above the average plus one standard deviation as peak signals. Furthermore, we determined that when the peak signal was larger than two pixels and smaller than ten pixels as one cluster. Finally, when the cluster of the two channels overlap with each other, we defined it as a colocalization event. We divided the colocalization events by the total clusters of PDLP1-mCherry to obtained the final colocalization percentage.

## Data availability

The data underlying this article are available in the article and in its online supplementary material.

## Acknowledgments

We gratefully thank Prof. Takayuki Kohchi, Prof. Ryuichi Nishihama, and Prof. Chang-Hsien Yang for the support of plasmids and materials. We also thank Dr. Barbara Kloeckener Gruissem for the critical review and editing of the manuscript.

## Author contributions

KJL: conceptualization; LTTT, SJT, HCL, CMH, and HYC: methodology; LTTT and SJT: formal analysis; LTTT, SJT, HCL, CMH, and HYC: investigation; KJL, LTTT, SJT, and HCL: data curation; KJL and LTTT: writing - original draft, review & editing; KJL, LTTT, SJT and HCL: visualization; KJL: supervision; KJL: funding acquisition

---

**Figure S1.**
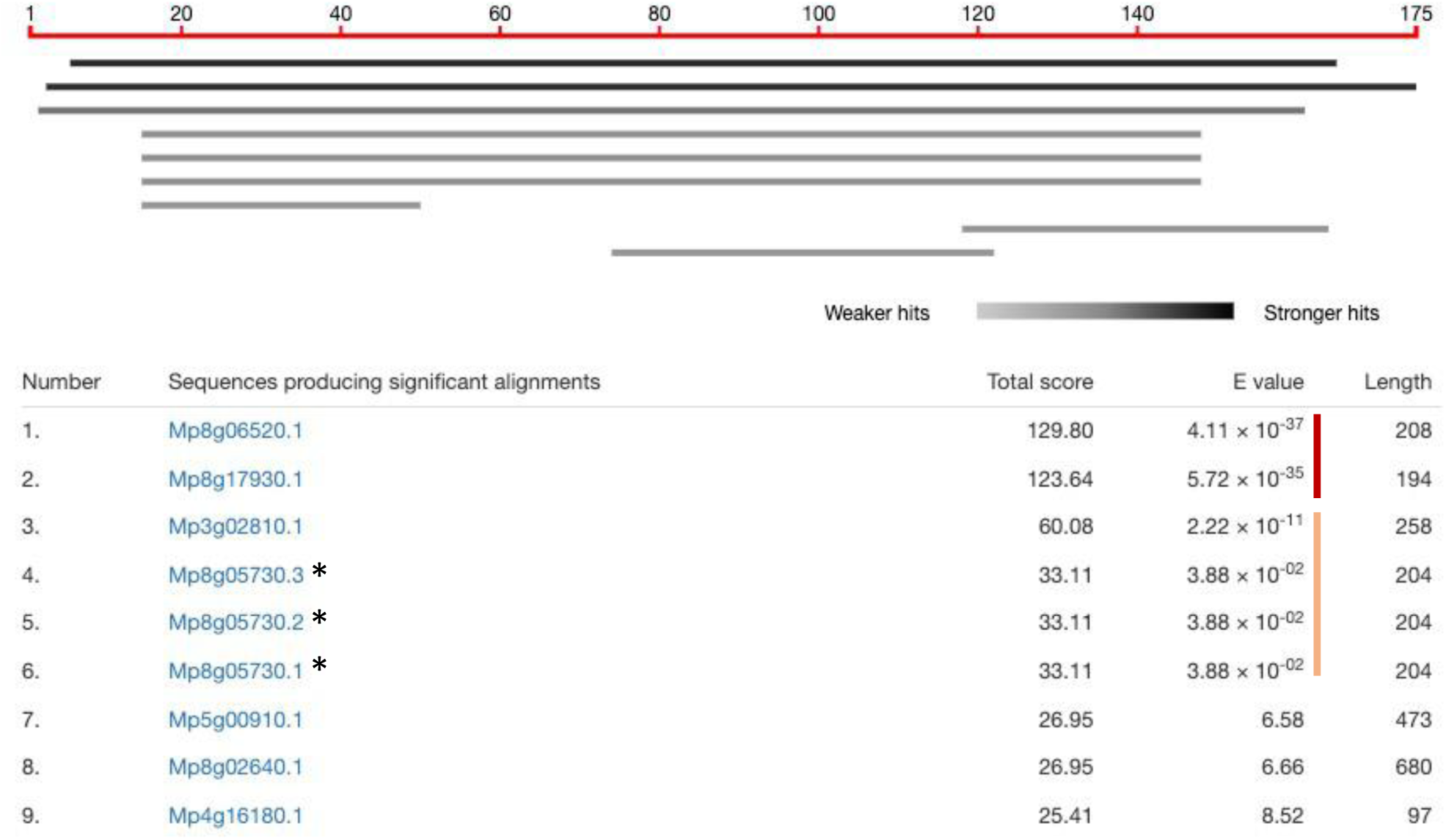
Identification and analysis of MpDUF1218 proteins. AtTVA was used as the query to perform BLASTP analysis on the Marpol base website (https://marchantia.info/). The red vertical line points to the two MpDUF1218 proteins with lower E-values, while the yellow vertical line points to the other two MpDUF1218 proteins with higher E-value. The proteins with lower E-values were chosen for further experiments. *Mp8g05730 underwent three revisions.

**Figure S2.**
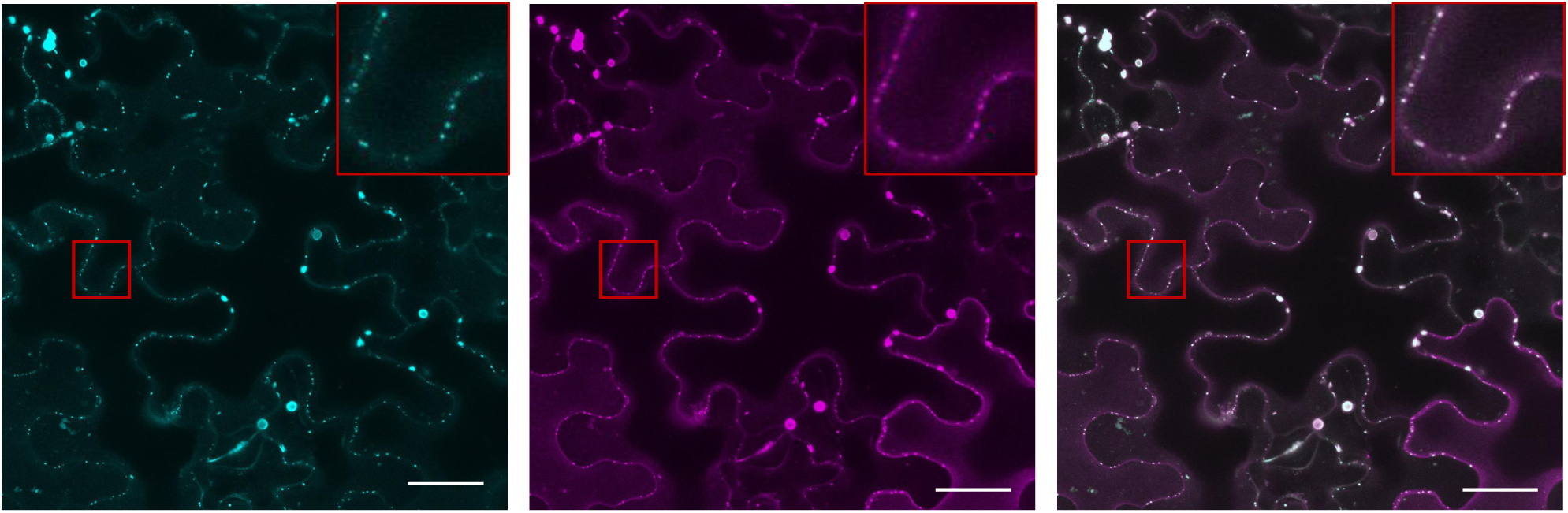
Colocalization of PDLP1-mCherry and aniline blue staining. (A, B) Fluorescent signals of aniline blue (A) and p35S::PDLP1-mCherry (B) expressed in the leaf epidermal cells of *N. benthamiana*. (C) Merged images of the previous two panels. Three-fold enlarged images of the red boxed sections are shown in the upper right corner of each panel. Scale bar, 30 μm.

**Figure S3.**
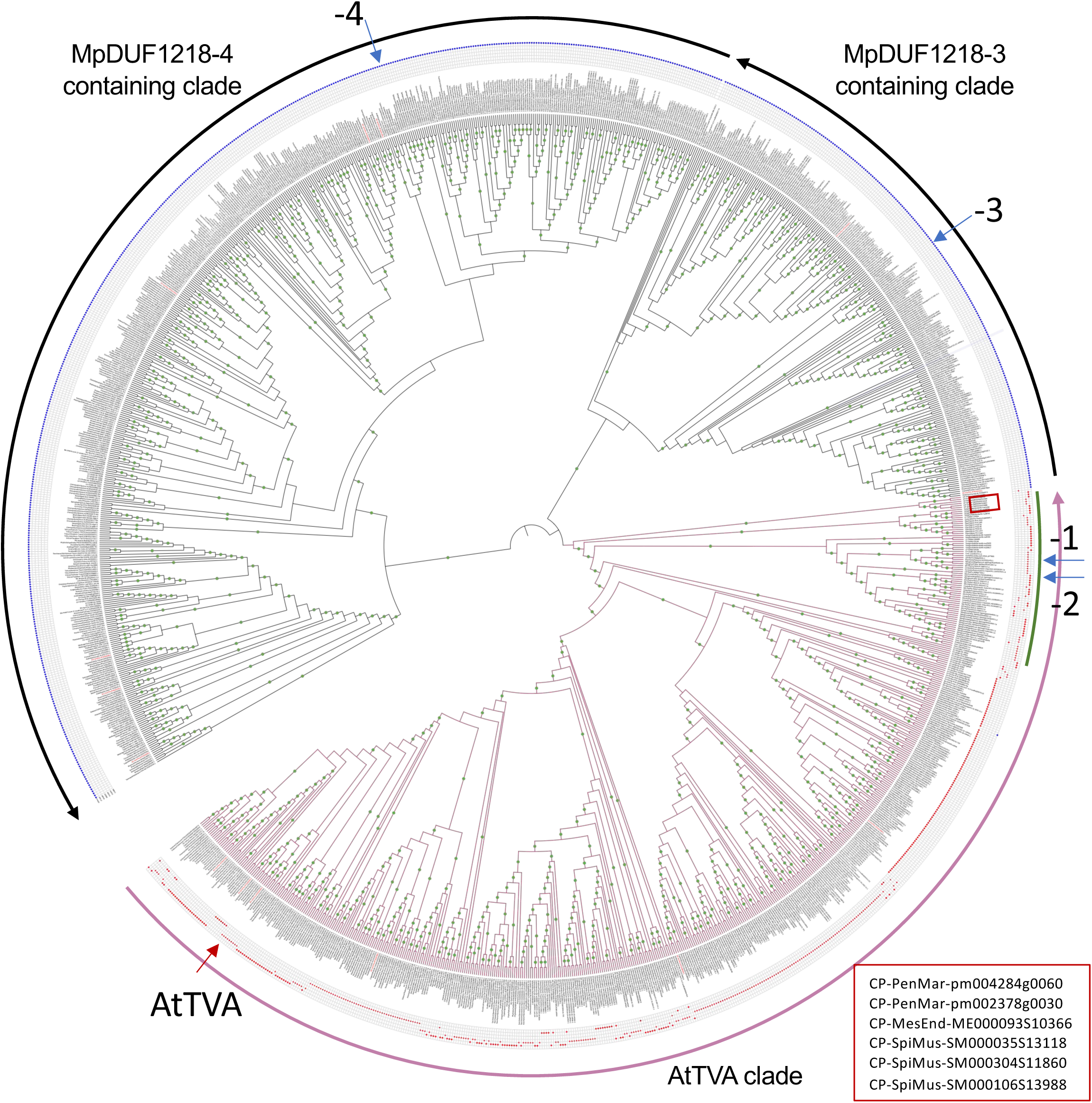
Phylogenetic tree of DUF1218 domain-containing proteins. The phylogenetic tree shows that DUF1218 domain-containing proteins can be identified in charophytes (shown in the red box and enlarged at the bottom right corner) and branched into AtTVA clade, MpDUF1218-3 containing clade, and MpDUF1218-4 containing clade. AtTVA clade contains MpDUF1218-1, -2, and six Arabidopsis DUF1218 domain proteins, indicated by the purple arrow at the outermost ring. The MpDUF1218-3 containing clade has 2 Arabidopsis DUF1218 domain proteins, indicated by the shorter black arrow at the outermost ring. The MpDUF1218-4 containing clade has another 6 Arabidopsis DUF1218 proteins, indicated by the longer black arrow at the outermost ring. MpDUF1218-1 to -4 are pointed by the blue arrows. AtTVA is pointed to by the red arrow. The green line shows the AtTVA-clade DUF1218 domain proteins identified from Bryophytes, Lycophytes, and Ferns. The red and blue dots in the middle ring indicate the closest Arabidopsis DUF1218 proteins by using the candidate protein sequence as queries to perform BLAST analysis. Green dots on the branches indicate synapomorphy. CP, Charophytes. PenMar, Penium *margaritaceum*. MesEnd, *Mesotaenium endlicherianum*. SpiMus, *Spiroglea muscicola*.

**Figure S4.**
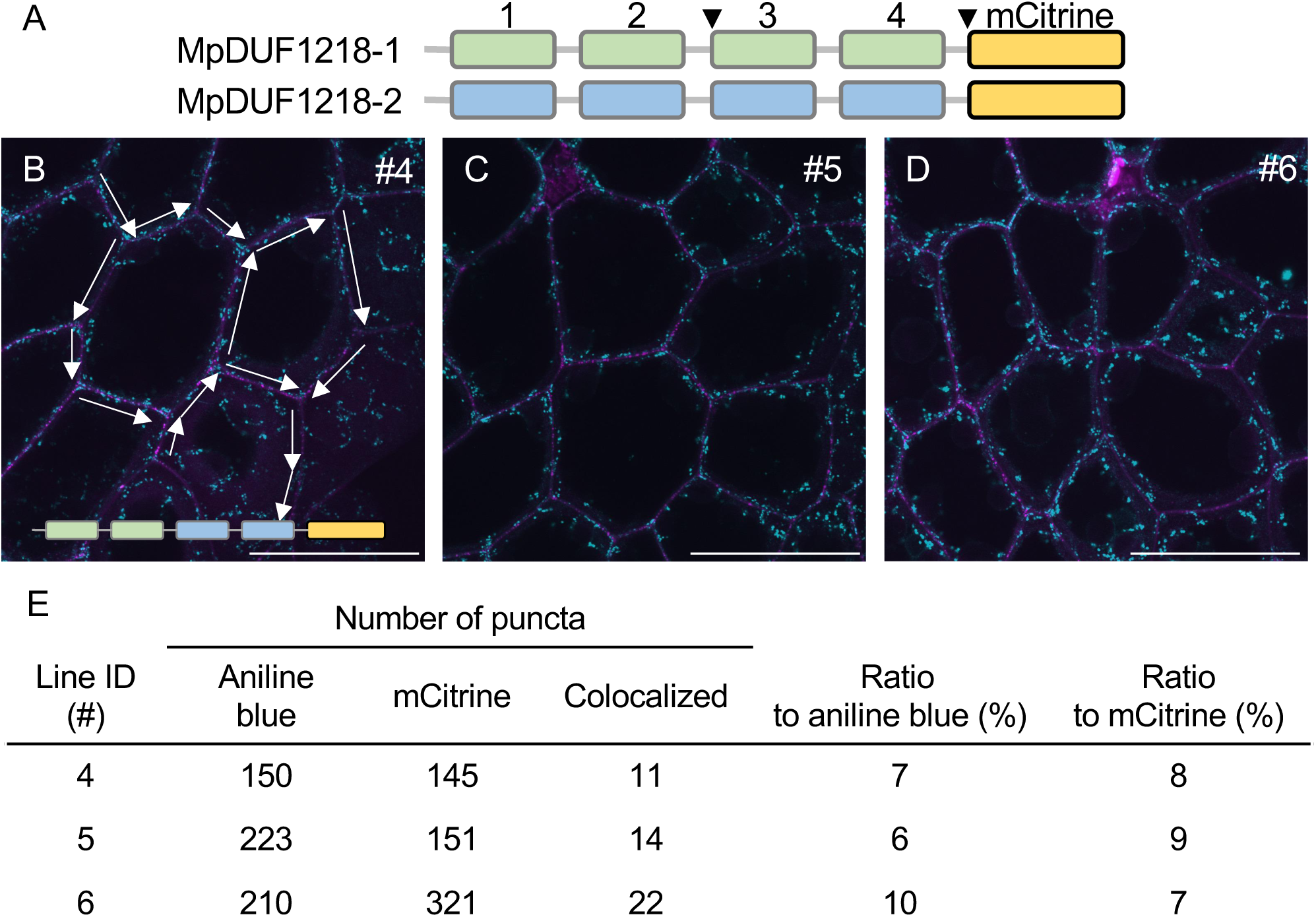
Colocalization analysis of the MpDUF1218-1 helix-swapped 3 and 4 with aniline blue staining in *M. polymorpha* 11-day-old gemmae. (A) Cartoon of the MpDUF1218 proteins. Blocks 1-4 represent helices (H) with the respective number above. Visual marker mCitrine (yellow box) was added to the C-terminus of all three proteins. When consecutive helices 3 and 4 were simultaneously swapped, the entire sequence between the arrowheads was swapped. (B-D) Fluorescent signals of the MpDUF1218-1 HS3,4 (helix-swapped 3 and 4) proteins in the epidermal cells of 11-day-old gemmae. The illustration of the helix-swapped protein is presented at the bottom of (B). Images were from three independent measurements of three independent transgenic lines. The line number of the transgenic plants is shown at the upper right corner of the image. The signals of MpDUF1218-1 HS3,4 are showing in cyan and the signals of aniline blue is showing in Magenta. Scale bar, 50 μm. White arrows in (B) illustrate the cell walls we selected for quantification. Only complete cell walls were selected for analysis. (E) Statistics of the puncta numbers in each image along selected cell walls. The calculation was done by Fiji (ImageJ, https://imagej.net/software/fiji/).

**Figure S5.**
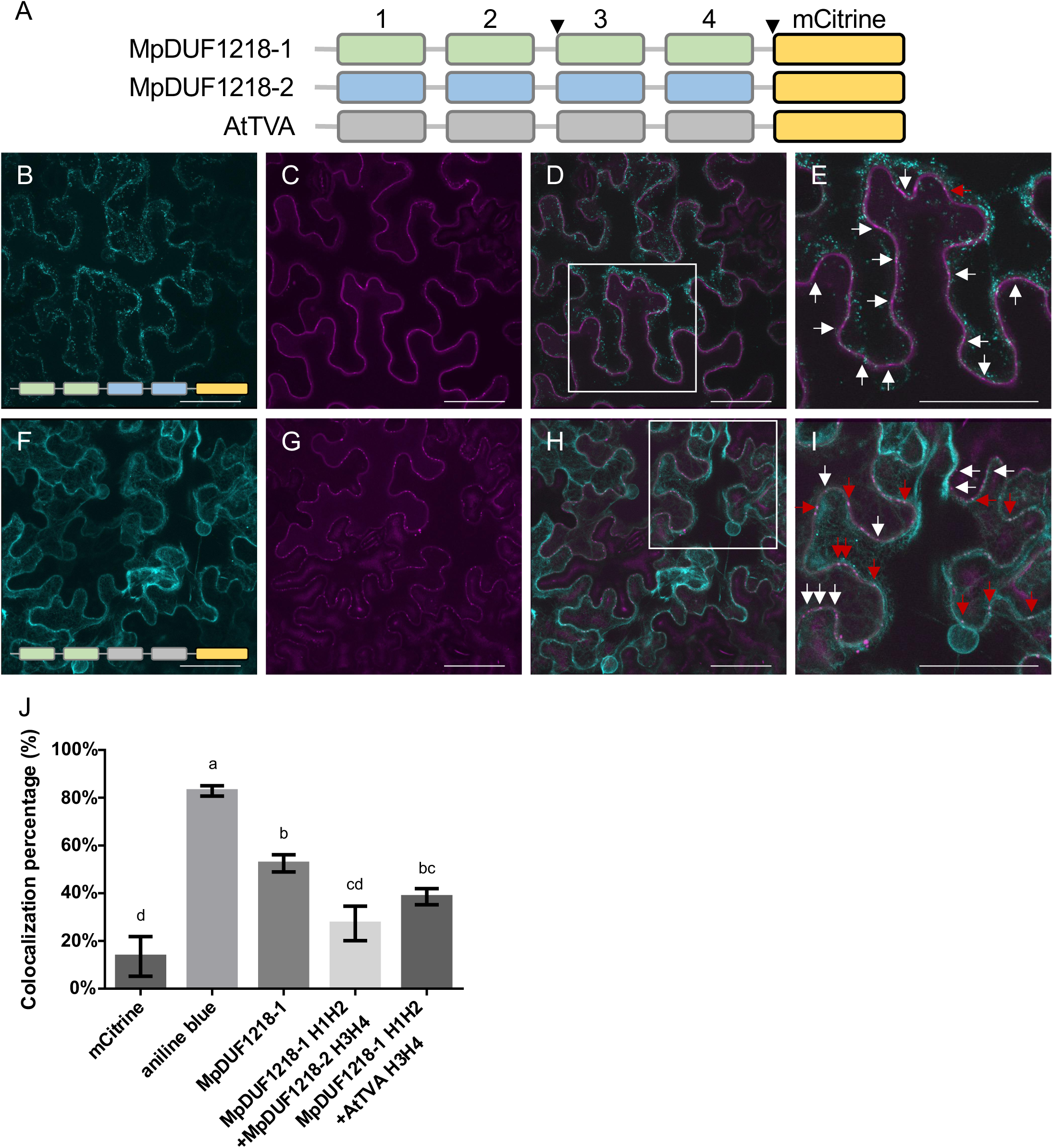
The subcellular localization of MpDUF1218-1 helix-swapped 3 and 4 with MpDUF1218-2 and AtTVA protein in *N. benthamiana*. (A) Cartoon of MpDUF1218-1, MpDUF1218-2 and AtTVA proteins. Blocks 1-4 represent helices showing the respective number above. Visual marker mCitrine (yellow box) was added to the C-terminus of all three proteins. When consecutive helices 3 and 4 were simultaneously swapped, the entire sequence between the arrowheads was swapped. (B, F) Fluorescent signals are presented from the indicated helix-swapped proteins at the bottom of each panel expressed in the leaf epidermal cells of *N. benthamiana*. (C, G) Plasmodesmata were visualized using p35S::PDLP1-mCherry expressed on the same leaf sections. (D, H) Merged images of the previous two panels, respectively. (E, I) Two-fold enlarged images from the red boxed sections in D and H. Red arrows point to the colocalization of the helix-swapped proteins with the plasmodesmata marker. White arrows point to the plasmodesmata markers only. In the merged images, the plasmodesmata marker alone showed magenta color while colocalized with the helix-swapped proteins and the plasmodesmata markers showed white color (D, E, H, I). Scale bar, 50 mm. (J) Colocalization analysis of the MpDUF1218-1 domain-swapped constructs with PDLP1-mCherry. mCitrine was used as the negative control while aniline blue staining was used as the positive control. Note that the same mCitrine and aniline blue data were reused as a comparison reference. n=3 for each construct. The data were analyzed by one-way ANOVA with Tukey’s HSD post hoc analysis.

**Figure S6.**
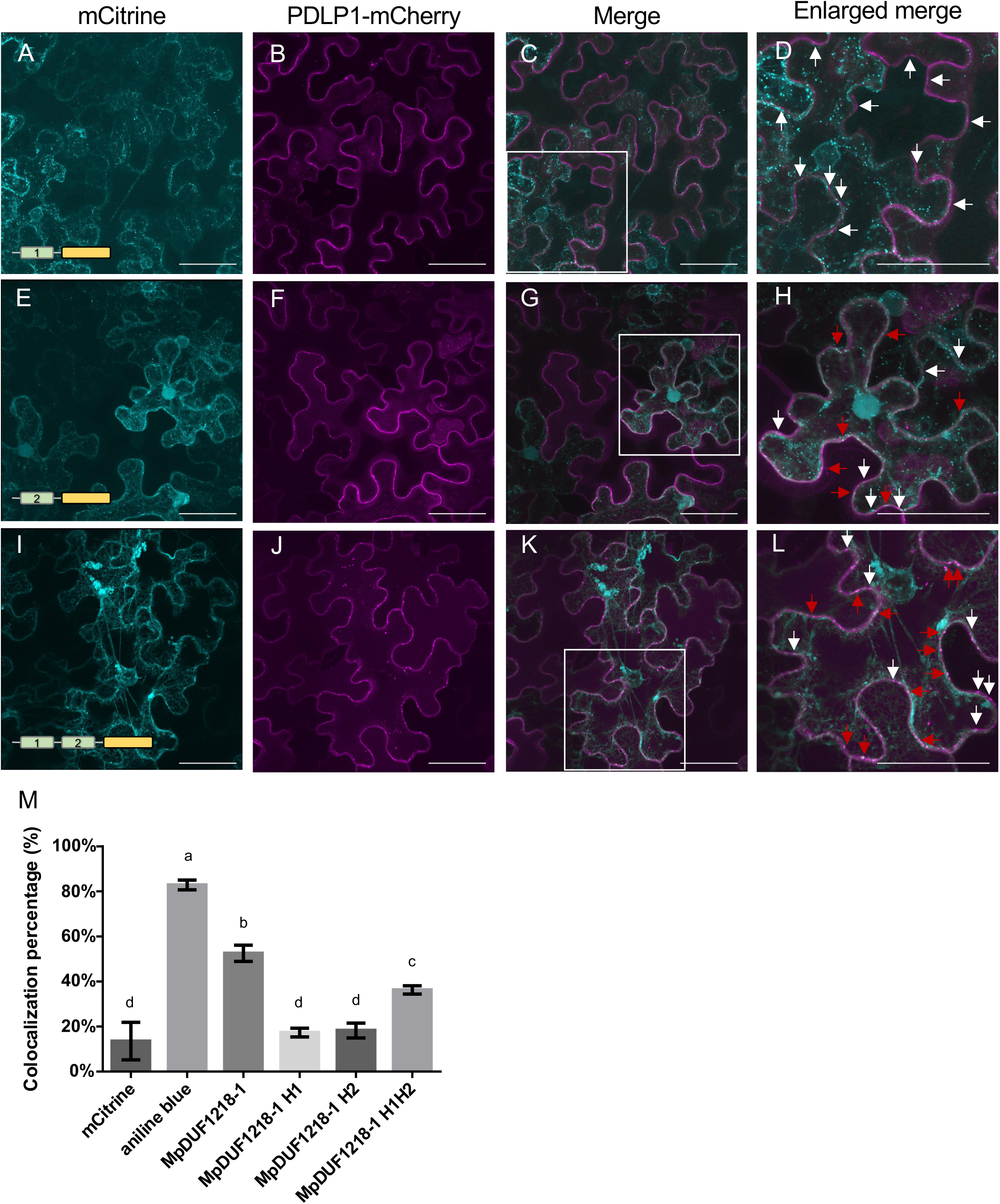
Helix1 and Helix2 of MpDUF1218-1 individually located at plasmodesmata in *N. benthamiana*, but not both helices together. (A, E, I) The fluorescent signals of the indicated fusion proteins expressed in the epidermal cells of *N. benthamiana*. (A) MpDUF1218-1 helix 1. (E) MpDUF1218-1 helix 2. (I) MpDUF1218-1 H1,2 (helix 1 and helix 2). (B, F, J) Plasmodesmata were visualized using p*35S*::PDLP1-mCherry expressed on the same leaf sections. (C, G, K) The merged images of the previous two panels, respectively. (D, H, L) Two-fold enlarged images of the red boxed sections in C, G, and K. In the merged image, plasmodesmata markers alone showed magenta color. When colocalized with the helix-deleted proteins, the plasmodesmata markers showed white color (D, H). Red arrows point to the colocalization of the candidate proteins with the plasmodesmata markers. White arrows point to the plasmodesmata markers without colocalization. Scale bar, 50 μm. (M) Colocalization analysis of the indicated MpDUF1218-1 helix constructs with PDLP1-mCherry. mCitrine was used as the negative control while aniline blue staining was used as the positive control. Note that the same mCitrine and aniline blue data were reused as a comparison reference. n=3 for each construct. The data were analyzed by one-way ANOVA with Tukey’s HSD post hoc analysis.

